# Structural characterization of a breast cancer-associated mutation in caveolin-1

**DOI:** 10.1101/2022.05.23.493104

**Authors:** Bing Han, Alican Gulsevin, Sarah Connolly, Ting Wang, Jason Porta, Ajit Tiwari, Angie Deng, Louise Chang, Yelena Peskova, Hassane S. Mchaoraub, Erkan Karakas, Melanie D. Ohi, Jens Meiler, Anne K. Kenworthy

**Affiliations:** Center for Membrane and Cell Physiology, University of Virginia, Charlottesville, VA USA; Department of Molecular Physiology and Biological Physics, University of Virginia School of Medicine, Charlottesville, VA, USA; Department of Chemistry, Vanderbilt University Nashville, TN, USA; Life Sciences Institute, University of Michigan, Ann Arbor, MI, USA; Department of Molecular Physiology and Biophysics, Vanderbilt University Nashville, TN, USA; Institute for Drug Discovery, Leipzig University, Germany; Department of Cell and Developmental Biology, University of Michigan School of Medicine, Ann Arbor, MI, USA

## Abstract

Caveolin-1 (CAV1) is a membrane sculpting protein that oligomerizes to generate flask-shaped invaginations of the plasma membrane known as caveolae. Mutations in CAV1 have been linked to multiple diseases in humans. Such mutations often interfere with oligomerization and the intracellular trafficking processes required for successful caveolae assembly, but the molecular mechanisms underlying these defects have not been structurally explained. Here, we investigate how a breast cancer-associated mutation in one of the most highly conserved residues in CAV1, P132L, affects CAV1 structure and oligomerization. We show that P132 is positioned at a major site of protomer-protomer interactions within the CAV1 complex, providing a structural explanation for why the mutant protein fails to homo-oligomerize correctly. Using a combination of computational, structural, biochemical, and cell biological approaches, we find that despite its homo-oligomerization defects P132L is capable of forming mixed hetero-oligomeric complexes with wild type CAV1 and that these complexes can be incorporated into caveolae. These findings provide insights into the fundamental mechanisms that control the formation of homo- and hetero-oligomers of caveolins that are essential for caveolae biogenesis, as well as how these processes are disrupted in human disease.

## INTRODUCTION

Flask-shaped invaginations of the plasma membrane known as caveolae function as critical regulators of human health and disease (1-3). The integral membrane protein caveolin-1 (CAV1) is a major structural component of caveolae and is required for caveolae formation in non-muscle cells (4). Its intrinsic ability to induce membrane curvature enables it to trigger the formation of caveolae-like structures even in bacteria (5,6). In mammals, CAV1 and caveolae are widely distributed in many tissues where they function in mechanosensation, lipid homeostasis, signaling, endocytosis, and mechanoprotection (1,7-9). Conversely, dysregulation of CAV1 and caveolae contribute to the development and progression of diseases such as cancer, asthma, pulmonary fibrosis, pulmonary arterial hypertension, chronic inflammatory respiratory diseases, and lipodystrophy (10-16).

Assembly of CAV1 into oligomeric complexes is an essential step in caveolae biogenesis, and defects in oligomerization can give rise to disease (17-21). Under normal conditions, CAV1 undergoes an initial oligomerization step that generates complexes ∼8S in size when analyzed by velocity gradient centrifugation, termed 8S complexes (20). 8S complexes then can form higher order 70S complexes that are ultimately incorporated into caveolae together with cavins and accessory proteins (20-24). Oligomerization of CAV1 depends on the presence of a region of the protein known as the oligomerization domain (18). Disruption of other regions of CAV1 can also interfere with its oligomerization, suggesting that the conformation of caveolins is highly optimized (19). One mutation known to interfere with oligomer formation is a proline (P) to leucine (L) mutation in one of the most highly conserved residues across caveolins, P132 (25,26). Identified in 16% of primary human breast cancers, P132L was subsequently shown to impact cell migration, invasion, and metastasis (27-29). While the prevalence of the P132L mutation in breast cancer was later questioned (30-33), an equivalent mutation in caveolin-3 (CAV3), P105L (denoted as P104L in many studies), gives rise to muscular dystrophies both in humans and model organisms (34-36). P132L thus represents a powerful tool to investigate how mutations in caveolins disrupt oligomerization and ultimately interfere with caveolae formation function.

CAV1 P132L exhibits a number of defects compared to its wild type counterpart. Instead of assembling into 8S oligomers, P132L forms a combination of monomers/dimers and high molecular weight complexes (19,20,37,38). It also is retained intracellularly (19,20,37), likely because of defects in oligomerization. A model for how P132L disrupts oligomer formation has been proposed (39) but was based on the behavior of a form of CAV1, which itself was unable to oligomerize fully. P132L has also been reported to disrupt the ability of WT CAV1 to traffic to the plasma membrane correctly (37). However, if and how P132L can directly oligomerize with WT CAV1 oligomers has yet to be directly tested.

Due to the lack of a high-resolution structure of CAV1, it has not been possible to study how disease-associated mutations in CAV1, such as P132L, disrupt the oligomerization and function of the protein. We have now determined a 3.5 Å cryo-EM structure of the human CAV1 8S complex (40). The complex consists of 11 CAV1 protomers organized into a disc with a central protruding β-barrel at the center (40). The structure uncovers extensive inter-protomer interactions that extend along the entire length of each protomer, locked in place by a previously unidentified pin motif close to the N-terminus and a C-terminal β-barrel extending from the cytoplasmic face of the disc (40).

In the current study, we use the structure as a framework for investigating the molecular basis for how the P132L mutation alters CAV1 structure and oligomerization states. We propose a new structure-based model predicting how the P132L mutation disrupts homo-oligomerization. We also provide evidence that P132L can form hetero-oligomeric complexes with WT CAV1 and that these complexes are capable of supporting caveolae biogenesis. These findings provide new insights into the mechanisms that control caveolae biogenesis at a molecular level and the structural impact of disease-associated mutants on this process. They may also help lead to the development of therapeutic tools for the treatment of disease.

## RESULTS

### P132 contributes to multiple protomer-protomer interfaces in the CAV1 complex

We first sought to understand how P132 contributes to the structure of 8S complexes. Based on the cryo-EM structure, P132 falls on α-helix 3 of the highly hydrophobic region of CAV1 termed the “intramembrane domain” (IMD). In the cryo-EM structure, the IMD corresponds to a predominantly helical structure lying along the plane parallel to the membrane surface (40). The positioning of the IMD in the complex suggests it fulfills two distinct functions: forming a flat membrane-facing surface and participating in numerous protomer-protomer interactions stabilizing the integrity of the complex. Within the IMD, P132 is located in a hydrophobic pocket between adjacent protomers, suggesting it primarily contributes to the packing of the complex rather than mediating membrane binding events (**Figure 1A-C**).

**Figure 1:**
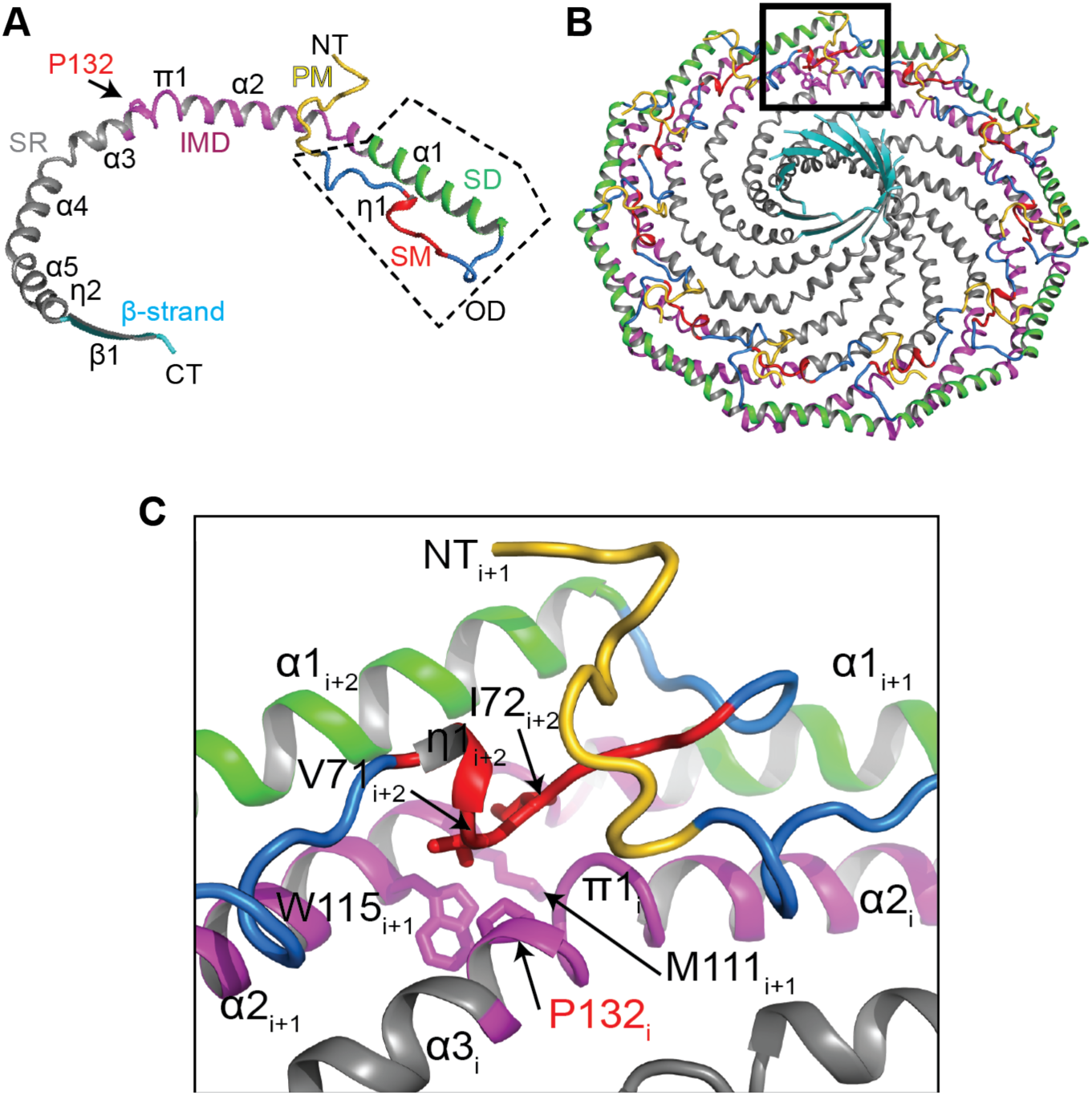
P132 is located at a major protomer-protomer interface in the CAV1 complex. **(A)** Structure of CAV1 protomer. The previously defined regions are labeled and colored: PM, pin motif (yellow); SM, signature motif (red); SD, scaffolding domain (green); and IMD, intermembrane domain (purple); SR, spoke region (grey); and β-strand (cyan). The oligomerization domain (OD), which contains the SM and SD, is indicated by the dashed box. P132 is shown in stick representation and indicated by an arrow. **(B)** The structure of the 8S CAV1 complex is shown in ribbon representation and colored as in panel A. **(C)** Zoomed up view of the boxed region in panel B. Select residues are shown in stick representation and labeled.

To probe the potential roles of P132 in supporting oligomerization, we examined its proximity to other protomers and conserved residues within this region. To facilitate this analysis, we refer to three adjacent protomers, *i, i+1*, and *i+2* (**Figure 1C**). The sidechain of P132 points into a hydrophobic pocket formed by M111 and the highly conserved W115 of the protomer *i+1* on the membrane-facing side and V71 and I72 of the protomer *i+2* on the cytoplasm-facing side. The latter two residues are located in the signature motif of CAV1, the most strongly conserved region of the protein across members of the caveolin gene family (41,42) (**Figure S1**). Thus, P132 makes important contacts with other conserved hydrophobic residues in several adjacent protomers within the complex. Furthermore, there is little space in the hydrophobic pocket to accommodate amino acids with larger side chains without structural rearrangements on the backbone atoms on the neighboring protomer. This suggests that mutations at this location of the structure could alter the ability of CAV1 protomers to tightly pack into a stable oligomer.

### Computational modeling predicts that the P132L mutation destabilizes CAV1 homo-oligomers

We next explored whether the P132L mutation alters the ability of CAV1 to homo-oligomerize. First, we used the cryo-EM 8S CAV1 structure as a template in Rosetta to ask to what extent P132 contributes to the stability of CAV1 oligomers. We computationally performed single amino acid substitutions at this position and calculated the corresponding ddG values relative to the P132P used as a control (**Table 1**). Positive ddG values indicate that the respective mutation destabilizes the structure, while negative ddG values indicate that the mutation is stabilizing. All amino acid substitutions at residue132 resulted in positive ddG values. However, the predicted extent of this destabilization varied depending on the specific amino acid substitution.

**Table 1.**
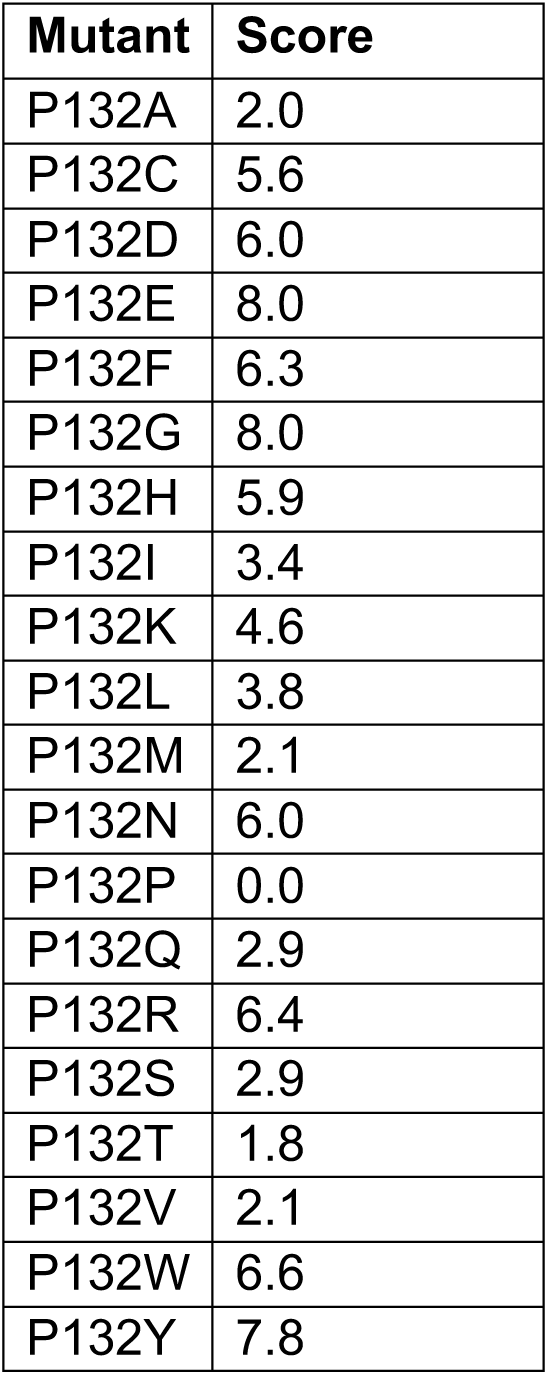
Impact on single amino acid substitutions at P132 on ddG values for a single CAV1 protomer. Rosetta ddG scores were calculated for single CAV1 protomers containing the indicated single amino acid substitutions at P132, relative to a P132P substitution and relaxed with 11-fold symmetry. Scores are reported in Rosetta Energy Units (REU).

| **Mutant** | **Score** |
| --- | --- |
| P132A | 2.0 |
| P132C | 5.6 |
| P132D | 6.0 |
| P132E | 8.0 |
| P132F | 6.3 |
| P132G | 8.0 |
| P132H | 5.9 |
| P132I | 3.4 |
| P132K | 4.6 |
| P132L | 3.8 |
| P132M | 2.1 |
| P132N | 6.0 |
| P132P | 0.0 |
| P132Q | 2.9 |
| P132R | 6.4 |
| P132S | 2.9 |
| P132T | 1.8 |
| P132V | 2.1 |
| P132W | 6.6 |
| P132Y | 7.8 |

For example, mutations such as P132T, P132M, and P132A are predicted to have relatively small effects, whereas P132G, P132E, and P132Y led to the highest ddG values indicating they are the most destabilizing (**Table 1**). P132L had a moderate effect on the stability of the system. Thus, while any amino acid substitution at position 132 is predicted to destabilize the oligomer, replacement of P132 with charged residues that would be buried in a hydrophobic environment, with bulky aromatic amino acids that can clash with other residues in the neighboring protomers, or with glycine that affects the helix backbone have particularly large destabilizing effects on the system. Overall, the Rosetta modeling suggests that the proline at position 132 fulfills a critical role for the stability of the CAV1 complex.

To explore the structural basis of the impact of the P132L mutation on the stability of CAV1 oligomers, we examined its effect on twelve neighboring residues that fall within 8 Å of this position (**Table 2**). This group includes four residues on either side of P132 on the same protomer (128-131 and 133-136 of chain *i*) and residues on adjacent protomers *i+2* (71, 72) and *i+1* (115) in the structure (**Figure 2A, B**). This approach focuses only on local changes and does not measure the impact of the P132L mutation on the global stability of the CAV1 complex, nor does it allow large-scale rearrangement of the CAV1 complex structure. Most residues within the vicinity of P132 had positive ddG values in response to the P132L mutation, suggesting this mutation is not favorable. Part of this effect was caused by the destabilization of the helix backbone due to the replacement of a restrained proline with a leucine residue as observed for the protomer *i* residues facing away from L132, and part of it was caused by the clashes induced by the bulkier leucine side chain that faced other hydrophobic residues in the vicinity. Bulky residues such as V71_*i+2*_, I72 _*i+2*_, and W115 _*i+1*_ showed the largest increase in ddG, which was also reflected by the movement of the V71_*i+2*_ and W115 _*i+1*_ sidechains away from the leucine residue in the mutant structure. These calculations provide further evidence that the P132L disrupts both the stability of the helix backbone and the hydrophobic packing of the residues in its vicinity, which in combination would be predicted to alter the ability of mutant protomers to back into an organized oligomeric complex.

**Table 2.** Effect of P132L mutation on adjacent residues. Per-residue Rosetta energies of twelve amino acids within 8 Å of P132L were subtracted from that of the P132P mutation. Residues predicted to be most strongly impacted by the P132L mutation are highlighted in bold. See **Figures 1 and 2** for the positioning of protomers *i, i+1*, and *i+2* in the complex. All ddG units are in Rosetta Energy Units (REU).

| ADJACENT RESIDUE | ddG |
| --- | --- |
| **115<sub>$i+1$</sub>** | **0.51** |
| **128<sub>$i$</sub>** | -0.09 |
| **129<sub>$i$</sub>** | 0.10 |
| **130<sub>$i$</sub>** | **0.55** |
| **131<sub>$i$</sub>** | 0.41 |
| **132<sub>$i$</sub>** | -0.21 |
| **133<sub>$i$</sub>** | 0.10 |
| **134<sub>$i$</sub>** | 0.01 |
| **135<sub>$i$</sub>** | -0.05 |
| **136<sub>$i$</sub>** | 0.22 |
| **71<sub>$i+2$</sub>** | **0.81** |
| **72<sub>$i+2$</sub>** | **0.74** |

**Table 3.** Effect of incorporation of increasing numbers of P132L mutants on the predicted stability of CAV1 hetero-oligomeric complexes. ddG was measured for CAV1 11-mers consisting of a mixture of the indicated number of copies of P132L together with WT CAV1. The total energy change of the complex and the energy change per number of P132Ls in the complex are reported as the score differences between P132L and P132P mutations. All energy numbers are in Rosetta Energy Units (REU).

| **# of copies of P132L** | **Total energy change (P132L - P132P)** | **Energy change per # of copies of P132L** |
| --- | --- | --- |
| 1 | 0.6 | 0.6 |
| 2 | 3.3 | 1.6 |
| 3 | 9.5 | 3.2 |
| 4 | 20.4 | 5.1 |
| 5 | 15.4 | 3.1 |
| 6 | 25.0 | 4.2 |
| 7 | 22.7 | 3.2 |
| 8 | 31.9 | 4.0 |
| 9 | 41.1 | 4.6 |
| 10 | 29.0 | 2.9 |
| 11 | 55.4 | 5.0 |

**Figure 2:**
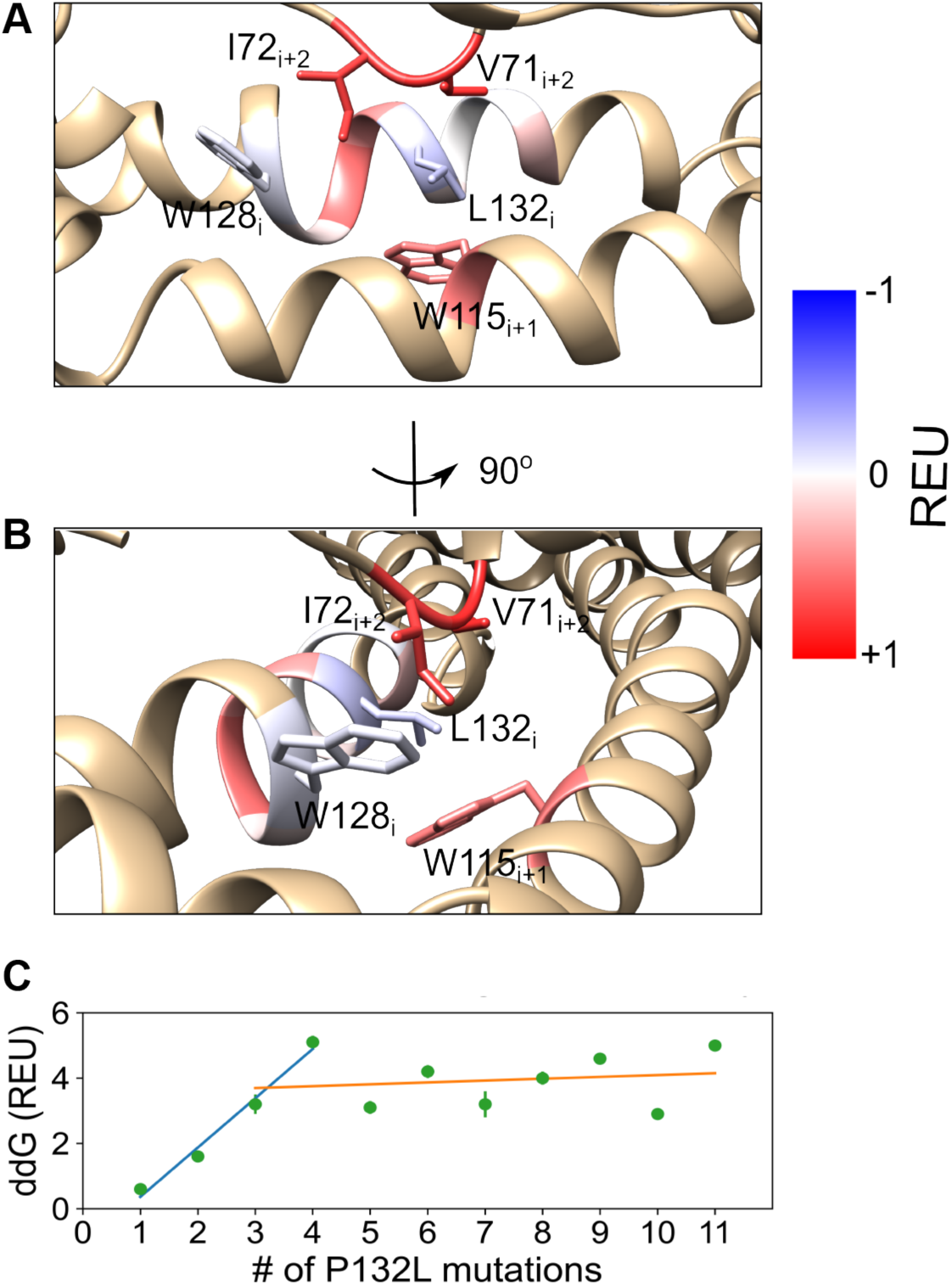
Impact of the P132L mutation on the stability of nearby residues and overall stability of CAV1 oligomers. (**A**) Rosetta ddG scores calculated for the residues within 8Å of L132 are color coded on the mutant structure. Blue indicates the P132L mutation has a stabilizing effect on the residue, and red indicates P132L’s effect is destabilizing. (**B**) As in A, except rotated 90 degrees. (**C**) Rosetta ddG changes per number of P132L mutations. Lines are drawn to guide the eye to highlight differences in predicted stability when 1-3 copies of P132L (blue line) versus 4 or more (orange line) are present in a mixed complex with WT CAV1. REU, Rosetta energy units.

### Computational modeling predicts P132L can form mixed complexes with WT CAV1

While it is currently unknown if P132L interacts directly with WT CAV1, results from cellular studies have suggested a model where P132L mutant protomers could oligomerize with WT CAV1 protomers. However, as our Rosetta modeling showed that P132L would likely disrupt homo-oligomerization, we hypothesized that the 8S CAV1 complexes potentially could only accommodate a limited number of P132L protomers before destabilizing the entire oligomer. This model predicts each P132L mutation would not only result in an increase in the ddG of the whole complex but also would increase the ddG calculated on a per-protomer basis. This is because each P132L mutation could destabilize its neighboring protomers in addition to the protomer that contains it. In other words, ddG per protomer would be expected to increase as a function of the increasing number of copies of mutant CAV1, rather than remaining fixed, as would be expected if the effects of the mutations were independent of one another.

To test this idea, we introduced the P132L mutation *in silico*, systematically increasing the number of mutant protomers from one to eleven while correspondingly decreasing the number of WT copies of CAV1. For simplicity, mutant protomers were introduced at neighboring positions within the complex. We then calculated the destabilization caused by these mutations as a function of the increasing number of P132L mutations in terms of Rosetta Energy Units (REU). A single P132L mutation destabilized the complex by ∼0.6 REU. This low value implies that the amount of destabilization is minor and may be compensated by the rest of the system (**Figure 2C**). However, as the number of copies of P132L was increased from one to four, the ddG calculated on a per-mutation basis increased drastically (**Figure 2C, blue line**). The per-protomer ddGs then level off for complexes containing five or more P132L mutations at a value indicative of substantial destabilization (**Figure 2C, orange line**). Overall, the introduction of eleven P132L mutations had a destabilizing effect of ∼55 REU, a value systematically higher than expected if the effects of the P132L mutations were additive rather than synergistic.

The Rosetta computational analyses show that P132 is an important residue located in a position where any amino acid substitution would diminish the stability of CAV1 homo-oligomers and limit the number of mutant protomers accommodated in hetero-oligomers. The effect of the P132L mutations is predicted to be synergistic, such that the increasing number of mutant protomers destabilize oligomers more strongly than if the effects of the mutations were independent of one another. Nevertheless, our results suggest that 8S complex formation may be possible if only a few copies of protomers bearing the P132L mutation are present in the complex.

### P132L and WT CAV1 co-assemble into 8S complexes in the heterologous E. coli model system

We next tested the effect of P132L on CAV1 8S complex formation using the heterologous *E. coli* model system. As we have shown, WT CAV1 expressed in *E. coli* assembles into 8S complexes that can be purified and analyzed biochemically and by electron microscopy (40,43). Furthermore, *E. coli*-expressed P132L has been reported to exhibit oligomerization defects similar to those seen in mammalian cells (5), suggesting *E. coli* is a good platform to investigate the effect of the P132L mutation on CAV1.

We first confirmed that P132L and WT CAV1 oligomerize as expected using blue native PAGE (BN-PAGE) to monitor their oligomerization status (19,38,43-45). For these experiments, we expressed both WT and P132L mutant CAV1 with and without fused Venus tag (CAV1, P132L, CAV1-mVenus, and P132L-mVenus) in *E. coli*. Following the cell lysis and purification of total membranes, we checked the oligomerization status of the proteins using blue native-PAGE followed by Western blotting. Two antibodies were used to specifically detect CAV1: an anti-N-term CAV1 antibody that recognizes both WT and P132L, and an anti-GFP antibody, which only detects the protein fused to mVenus. In blue native gels, WT CAV1-mVenus migrated as a high molecular weight band in a position expected for 8S complexes (**Figure 3**) (43). In contrast, P132L failed to migrate as a high molecular weight band and instead migrated as a smear (**Figure 3**), consistent with its reported oligomerization defects (5). Strikingly, however, P132L-mVenus co-migrated with WT CAV1 when both proteins were co-expressed (**Figure 3**, lane 5, red brackets and arrows). These results show that P132L CAV1 can form mixed 8S complexes with WT-CAV1 when expressed in *E. coli* as predicted by our Rosetta calculations. On the other hand, none of the high molecular bands were disrupted by co-expressing with P132L, indicating that the incorporation of P132L does not destabilize these complexes (**Figure 3**, lane 5 and 8).

**Figure 3:**
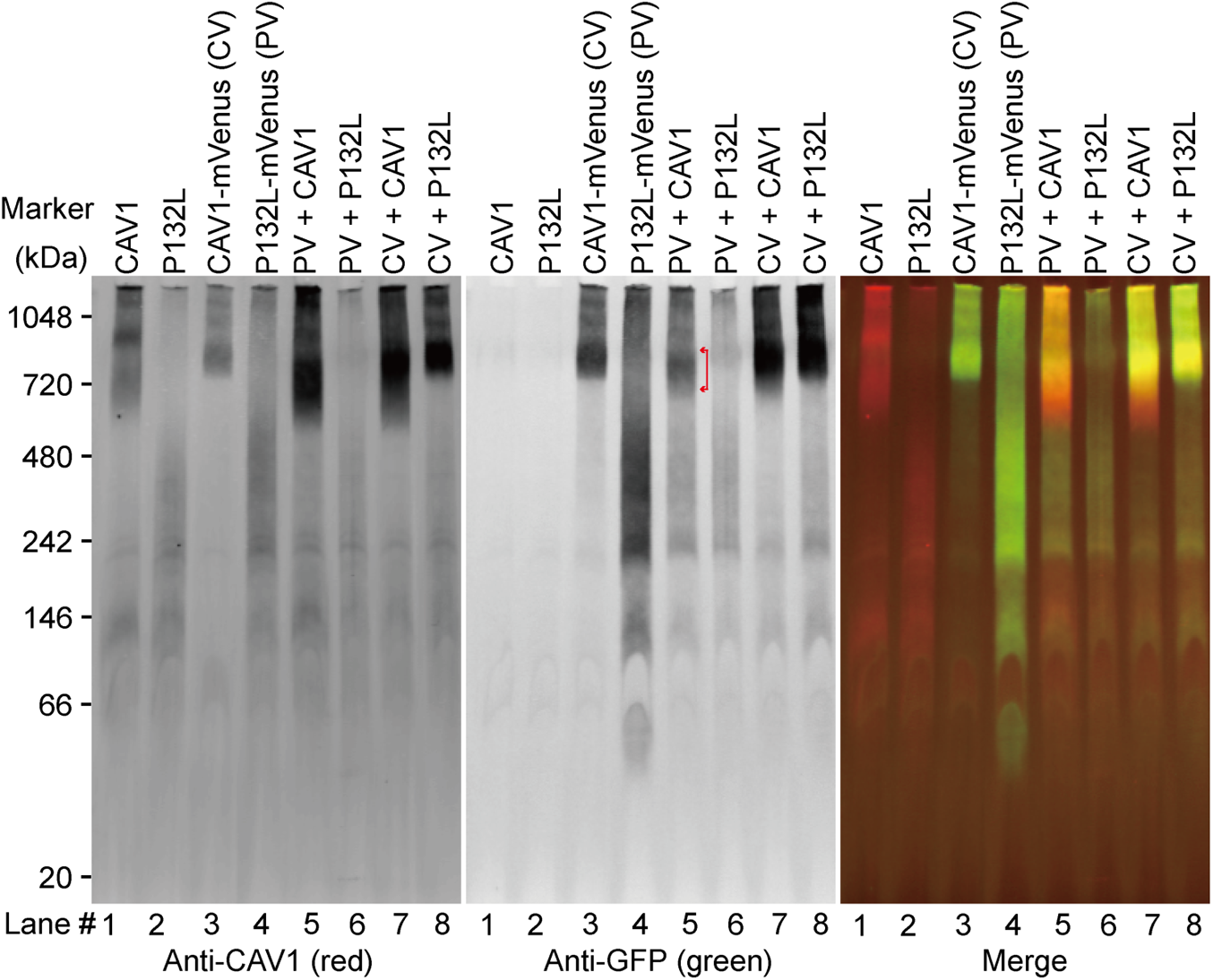
P132L forms mixed 8S oligomers with WT CAV1 in *E. coli* cells as assessed by BN-PAGE. Total membrane from *E. coli* cells expressing the indicated constructs were lysed in C12M and subjected to BN-PAGE followed by western blotting for mVenus (GFP, green) and CAV1 (red). Lane numbers were labeled at the bottom. The range of high molecular bands is indicated with red brackets and arrows in middle panel. PV, P132L-Venus; CV, CAV1-Venus.

Next, we examined the organization of the oligomers formed by P132L and the P132L/ WT CAV1 mixed complexes using negative stain electron microscopy. For these experiments, WT CAV1, CAV1-mVenus, P132L, and P132L mVenus were expressed singly or in combination in *E. coli*, purified using nickel beads, and fractionated using FPLC (**Figure S2**). Pooled samples were then negatively stained and imaged by EM.

Consistent with our previous results (43), CAV1-mVenus or CAV1 fractionated into two peaks, P1 and P2 (**Figure S2A**,**C**). We focused our analysis on the P1 fractions, which consist of 8S complexes (43). These complexes are disc shaped, ∼15 nm in diameter, and contain a central protrusion when viewed from the side (**Figure 4A,B**, **Figure S3A**,**B**). Examples of both *en face* views and side views of the disc-shaped 8S complexes can be seen in 2D average classes of CAV1 and CAV1-mVenus (**Figure 4A,B**, **Figure S3A**,**B**). CAV1 oligomers can also form 8S complex “dimers” composed of two 8S complexes interacting with each other via their central protrusions **(Figure 4B,I**). CAV1-mVenus particles contain an additional fan-shaped density emanating from the central protrusion visible in the side views consistent with the C-terminal mVenus tag (**Figure 4A,J**). Unlike CAV1, CAV1-mVenus complexes do not form dimers (**Figure 4A**).

**Figure 4:**
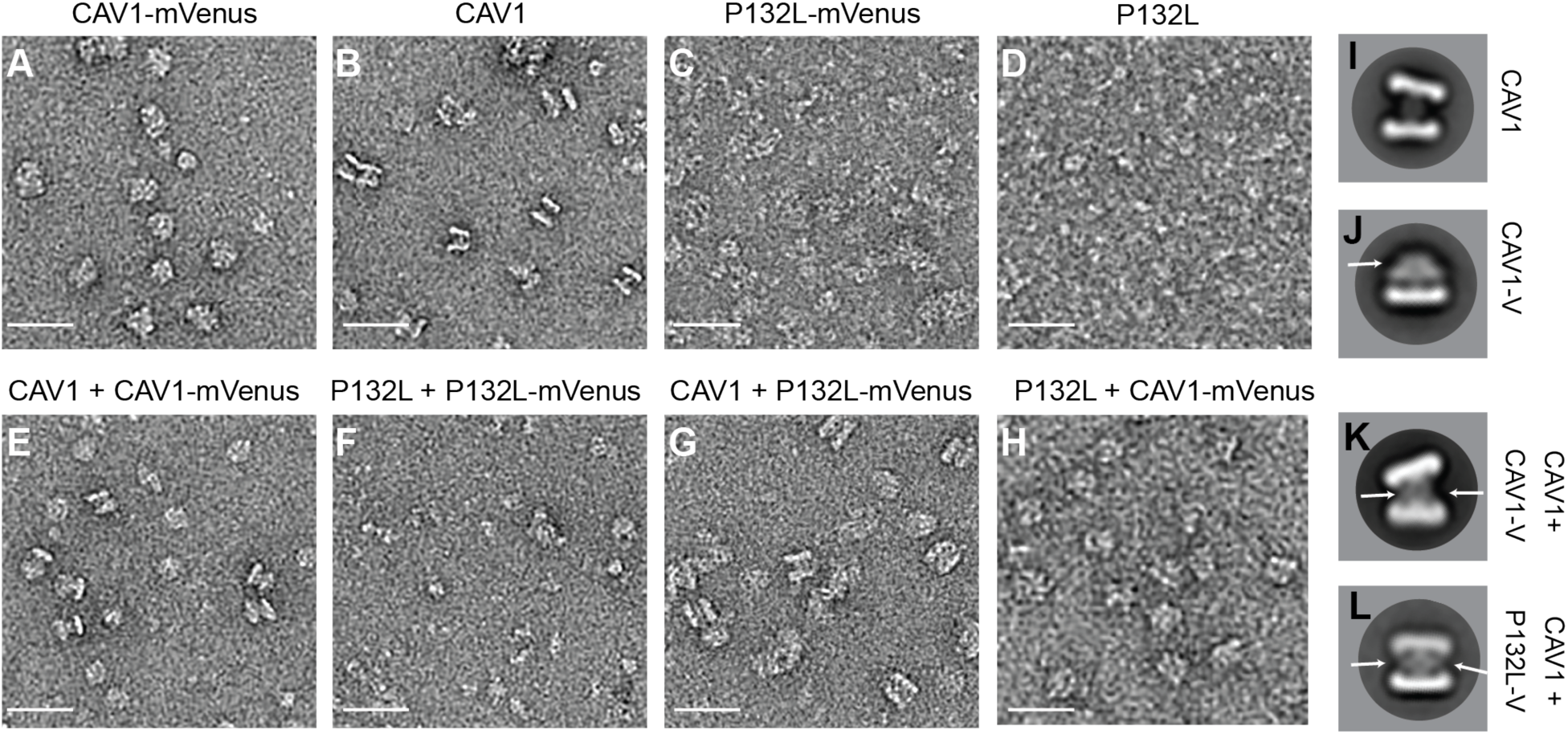
P132L forms mixed 8S complexes with WT CAV1 as assessed by negative stain electron microscopy. Purified CAV1 complexes were negatively stained and visualized by electron microscopy. Representative images of negatively stained P1 fractions of (**A**) CAV1-mVenus, (**B**) CAV1, (**C**) P132L-Venus, (**D**) P132L, (**E**) CAV1 + CAV1-mVenus, (**F**) P132L + P132L-mVenus, (**G**) CAV1 + P132L-mVenus, and (**H**) P132L + CAV1-mVenus. Scale bar, 30 nm. Representative 2D averages of (**I**) CAV1 complexes, (**J**) CAV1-mVenus 8S complexes, (**K**) CAV1 + CAV1-mVenus mixed complexes, and (**L**) CAV1 + P132L-mVenus mixed complexes. Arrow marks regions of additional density from the Venus tag. Scale bars, 30 nm.

We then analyzed the structure of oligomeric complexes formed by P132L and P132L-mVenus. P132L displayed a more complicated FPLC profile than CAV1 complexes, consisting of at least seven shoulders containing heterogeneous particles when examined by negative stain EM (**Figures 4C**, **S2B**,**D**). The P132L and P132L-mVenus complexes in the P1 fraction were not structurally well organized and thus not amenable for 2D averaging (**Figure 4C,D**).

Next, we visualized mixed complexes formed by co-expression of WT-CAV1 and P132L-mVenus. We tagged the mutant with the Venus tag because it contributes an extra fan-like density that can be readily observed in the side views of the complexes (**Figure 4J**). We reasoned that the mVenus-tag should contribute extra density in the central β-barrel region if WT CAV1 and P132L-mVenus form hetero-oligomeric complexes. As a control, we first examined complexes purified from *E. coli* co-expressing WT CAV1 and WT CAV1-mVenus (**Figure 4E**). 2D averages of the side views of 8S CAV1 dimers had increased central density in the stalk region when compared to CAV1 complexes alone (**Figure 4I,K**), showing that the mVenus tag can be used to identify mixed complexes. Next, we examined 8S complexes purified from cells co-expressing WT CAV1 and P132L-mVenus. As predicted by our computational model and blue native gel analysis, the 2D class averages of dimeric complexes have additional densities in the central region similar to our control experiment co-expressing WT CAV1 and WT CAV1-mVenus (**Figure 4L**, white arrows). The diameter of the discs visible in the side views of mixed complexes formed between CAV1 + P132L-mVenus was comparable to those formed by WT CAV1 + WT CAV1-mVenus. These findings suggest that P132L and WT CAV1 are indeed capable of co-assembling into 8S complexes similar in size to those generated by WT CAV1.

### P132L and WT CAV1 form mixed detergent-insoluble 8S complexes in mammalian cells

We next asked whether P132L and WT CAV1 can co-assemble into 8S-like complexes in mammalian cells. To test this, we expressed WT CAV1 and P132L constructs individually or together in cells lacking endogenous caveolin expression, *CAV1*^-/-^ mouse embryonic fibroblast (MEF) cells. Previous studies have established these cells are a good model to reconstitute steps in caveolae assembly (26,44,45). After transfection with CAV1 constructs, the cells were allowed to express the protein for 16 h and then lysed with 0.5% TX-100. The lysates were then subjected to sucrose density centrifugation (20). WT CAV1 is enriched in 2 peaks corresponding to the position of 8S and 70S complexes (**Figure 5A**). P132L was mostly fractionated as low molecular weight species (**Figure 5B**), but was recruited into both 8S and 70S complexes when co-expressed with WT CAV1 **(Figure 5C)**. Furthermore, the stability of the mixed 8S complexes was similar to those formed from WT CAV1 alone, as evidenced by their resistance to lysis in 0.4% SDS plus TX-100 (**Figure 5D-F**).

**Figure 5:**
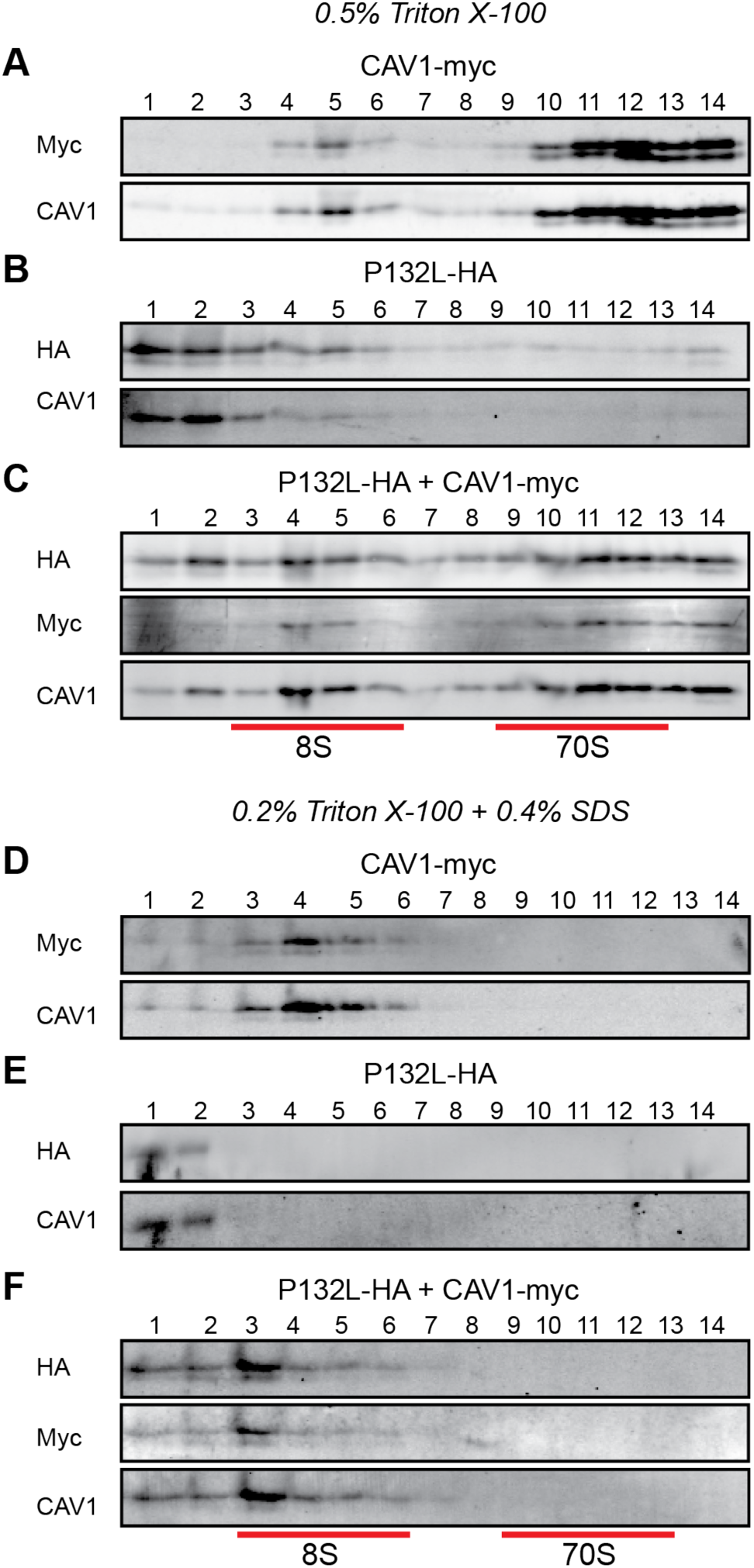
P132L forms mixed 8S and 70S complexes with WT CAV1 in CAV1 ^-/-^ MEF cells. MEF CAV1 ^-/-^ cells transiently transfected with **(A, D)** CAV1-Myc, **(B, E)** P132L-HA or **(C, F)** CAV1-Myc plus P132L-HA were lysed in either 0.5% Triton-X-100 **(A-C)** or 0.2% Triton X-100 plus 0.4% SDS **(D-F)** at room temperature. Extracts were run through 10-40% sucrose velocity gradients and fractions were analyzed by SDS-PAGE/western blot. Fraction numbers are indicated at the top of the blots and the positions of 8S and 70S complexes are indicated by red lines.

Another characteristic biochemical feature of WT CAV1 is that it associates with buoyant detergent-resistant membranes (DRMs) when properly incorporated into caveolae (19,44-46). In contrast, P132L is excluded from DRMs (37,38). We thus wondered whether the mixed complexes formed by WT CAV1 and P132L are detergent resistant. To address this question, MEF *CAV1* ^-/-^ cells overexpressing WT-CAV1 and P132L were extracted with 0.5% cold Triton X-100 followed by sucrose density fractionation (44,45). WT CAV1 was associated primarily with DRM fractions, as expected (**Figure 6A**). Although P132L localized to detergent soluble fractions when expressed on its own **(Figure 6B**), it shifted to the DRM fractions upon co-expression with WT CAV1 (**Figure 6C**). Thus, P132L can become incorporated together with WT CAV1 into detergent resistant 8S complexes in mammalian cells.

**Figure 6:**
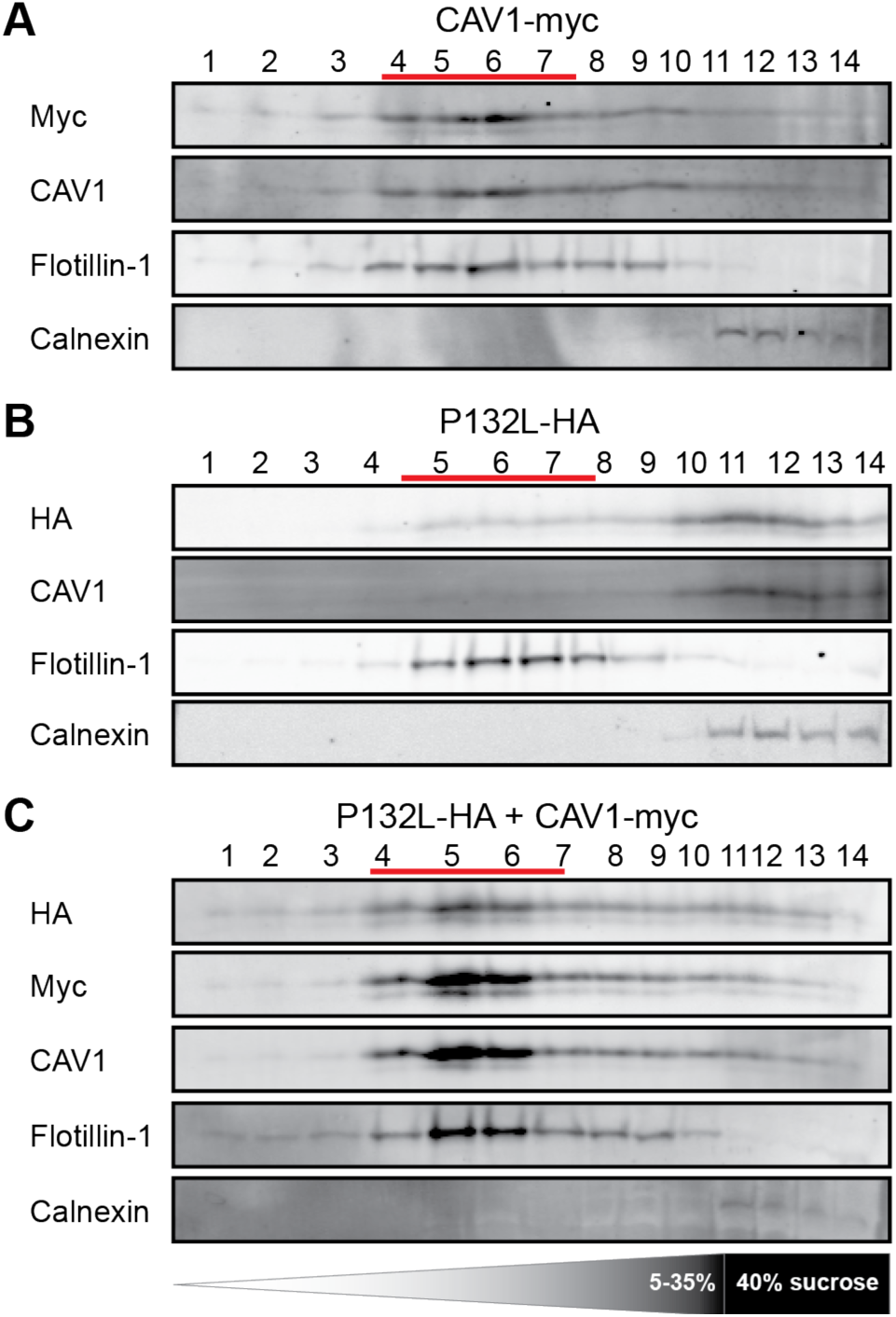
P132L is recruited into DRMs by WT CAV1 in CAV1^-/-^ MEFs. DRMs were isolated from CAV1 ^-/-^ MEF cells transiently transfected with **(A)** CAV1-Myc, **(B)** P132L-HA or **(C)** CAV1-Myc plus P132L-HA. Fractions were analyzed by SDS-PAGE/western blotting. Flotillin and calnexin were used as markers for DRMs and detergent soluble fractions, respectively. Fraction numbers are indicated at the top of the blots and the positions of the DRM fractions are indicated by red lines.

### P132L and WT CAV1 co-assemble into caveolae in mammalian cells

8S and 70S complexes function as the fundamental building blocks of caveolae (20). We, therefore, wondered whether mixed 8S complexes formed by P132L and WT CAV1 could support caveolae biogenesis. To address this question, we utilized *CAV1* ^-/-^ MEFs to assay for caveolae formation. Reconstitution of caveolae formation in these cells can be assessed by monitoring recruitment of Cavin-1 to puncta induced by exogenous expression of CAV1 as a readout of successful caveolae assembly (26,44,45).

We first confirmed that the expression of WT CAV1 in CAV1^-/-^ MEFs recruits endogenous cavin-1 to the plasma membrane, where it colocalizes with caveolin-positive puncta as visualized by AiryScan confocal microscopy (**Figure 7A**) or TIRF microscopy **(Figure S4A)**. In contrast, P132L localized primarily to the Golgi complex, and little P132L or Cavin-1 staining could be detected at the plasma membrane (**Figure 7B, S4B**). We then asked whether WT CAV1 can recruit P132L into caveolae upon co-expression of the two proteins. In this analysis, P132L colocalized extensively with WT CAV1 and cavin-1 in puncta at the cell surface (**Figure 7C, S4C**).

**Figure 7:**
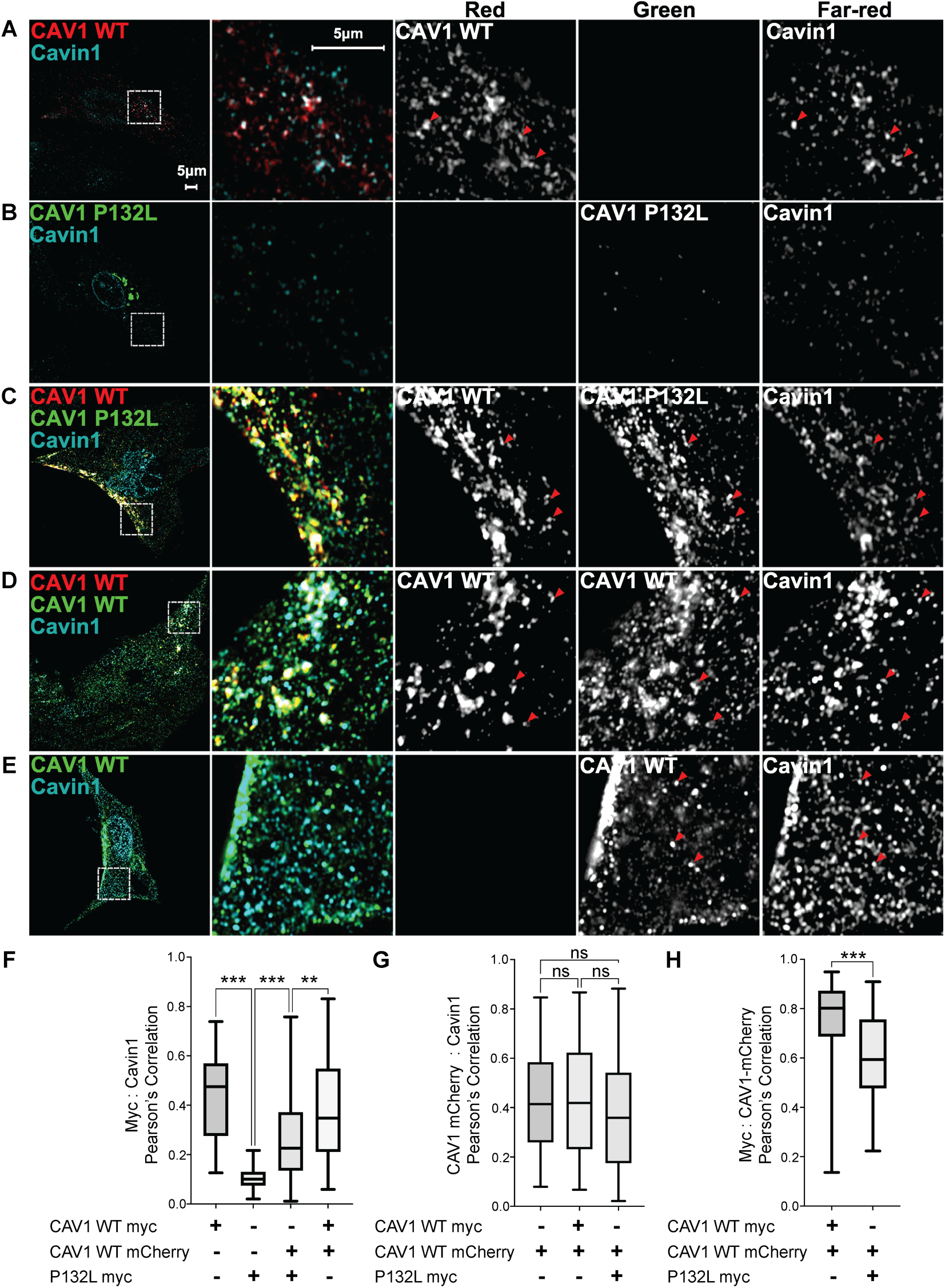
P132L localization defect is rescued by co-expression of WT CAV1 in CAV1 ^-/-^ MEFs. Representative AiryScan images are shown for CAV1 ^-/-^ MEF cells expressing **(A)** CAV1-mCherry, **(B)** P132L-Myc, **(C)**, CAV1-mCherry plus P132L-Myc **(D)** CAV1-mCherry plus CAV1-Myc or **(E)** CAV1-Myc. Cells were allowed to express the indicated constructs for 24h, fixed, and immunostained for endogenous cavin and myc-tagged constructs prior to imaging. Bar, 5 !m. In A-E, red arrowheads point to examples of colocalized puncta. **(F-H)** Pearson’s correlation analysis (n = 90 ROIs from 3 independent experiments). n.s., not significant. A one-way ANOVA with Tukey’s test (≥3 groups) or an unpaired Student’s t test (2 groups) was used to calculate P-values. N.s., not significant.

In addition to the cavins, caveolae incorporate accessory proteins that help regulate their morphology and dynamics (24,47-51). We tested whether two accessory proteins, PACSIN2 and EHD2, are found in mixed caveolae containing P132L. EHD2 and PACSIN2 are partially colocalized with caveolae containing either WT CAV1 complexes or P132L/WT complexes (**Figures S5, S6**). We conclude that mixed 8S complexes formed by P132L and WT CAV1 can successfully assemble into caveolae that contain a normal complement of accessory proteins.

## Discussion

Our results suggest the following model for how P132L disrupts caveolae assembly: We propose that P132 normally contributes to hydrophobic packing between protomers along the outer ring of the 8S complex. The introduction of a leucine residue at P132 introduces additional bulk facing the residues in neighboring protomers. This causes clashes with other conserved hydrophobic residues of CAV1 and prevents homo-oligomers containing more than a few copies of P132L from forming. Despite its homo-oligomerization defects, P132L can form hetero-oligomers with WT CAV1. Hetero-oligomers containing greater than 3 copies of P132L are predicted to be unstable using Rosetta calculations. Such irregular complexes would likely be rapidly degraded. In contrast, our computational analysis suggests that up to three copies of P132L can be incorporated into 8S complexes with WT CAV1 without compromising the integrity of the complex. These complexes can be detected experimentally using multiple techniques, appear structurally similar to those formed exclusively by WT CAV1, and can even become incorporated into caveolae that recruit caveolae accessory proteins appropriately. Taken together, these findings suggest that P132L expression could potentially interfere with the normal function of CAV1 in several ways. For example, mixed caveolae formed by co-expression of P132L and WT CAV1 might fail to support one or more normal functions of caveolae. Alternatively, intracellular pools of incompletely oligomerized P132L could potentially interfere with the normal functions of caveolin by competing for critical binding partners of CAV1, disrupting caveolae-dependent signaling or trafficking pathways, and/or inducing proteostatic stress.

Several of our current findings contradict previous reports. Based on structural analysis of CAV1 within the context of a complete 8S complex, we show that P132L primarily disrupts hydrophobic packing. This conflicts with previous work based on studies of a truncated form of CAV1 (residues 62-178) that suggested the P132L mutation alters the secondary structure of the IMD by extending an α-helix (39). We speculate the truncated form of the protein, which behaved as a monomer, may have undergone structural transitions that do not occur in the context of the fully assembled oligomer.

Our results do agree, however, with the conclusion that no other hydrophobic residue can substitute for the function of P132 in this position (39). We also found that P132L can become incorporated into caveolae together with WT CAV1. This finding was surprising in light of previous reports that expression of P132L cause WT CAV1 to become trapped intracellularly (37). However, these original studies were carried out in cells which we showed to have a tendency to accumulate overexpressed CAV1 in the perinuclear region as a consequence of aggresome formation (52,53). On the other hand, stable exogenous expression of P132L has also been reported to have no effect on caveolae formation in H1299 cells, a cell line expressing endogenous CAV1 (28).

Taken together, these findings highlight the complexities of studying caveolins and the importance of studying the proteins under conditions that preserve their ability to oligomerize and assemble into caveolae.

In addition to P132L, a variety of other pathogenic mutations in caveolins have been identified in humans (12,44,45,54-63). The most direct equivalent to P132L is a P105L mutation in CAV3 associated with muscular dystrophy (56,57). Muscle biopsies from patients harboring the P105L mutation with an autosomal form of dominant limb-girdle muscular dystrophy contain considerably decreased CAV3 levels compared to controls (56). Like CAV1 P132L, in mammalian heterologous expression systems, the P105L mutant protein accumulates in the perinuclear region and can trap WT CAV3 intracellularly (35,64). However, the nature of the oligomerization defects appears to be somewhat different, as P105L CAV3 tended to form much larger oligomers than WT CAV3 (35). Although a high-resolution structure of CAV3 has yet to be determined, it seems likely that the P105L mutation plays a similar role in destabilizing the structure of CAV3 complexes, as is the case for CAV1. To this point, it was recently suggested that P105 contributes to hydrophobic packing based on the modeling of the structure of CAV3 (65).

In contrast to P105L CAV3, disease-associated frameshift mutations in the C-terminal region of CAV1 appear to operate by a different mechanism than P132L. A mutation identified in patients with both familial and idiopathic forms of PAH, P158P, gives rise to a new C-terminus that is one residue longer than WT CAV1 and introduces a de novo ER retention signal (45,66). A different frameshift mutation, F160X, was found in a patient with both PAH and CGL and leads to premature termination of the protein (44,58,59). Unlike P132L, F160X can form 8S-like complexes and even support caveolae assembly when expressed in CAV1^-/-^ MEFs (44). However, mixed complexes formed by WT CAV1 and F160X in patient cells are destabilized compared to those formed by WT CAV1 (44). This destabilization likely results from the inability of F160X to form the central β-barrel of the 8S complex, evidenced by the loss of the central protrusion in these complexes when observed by negative stain EM (43). Both P158P and F160X can also form 8S-like complexes in *E. coli* (43). While disc shaped, these complexes are less regular than WT 8S oligomers, suggesting that disruption of the C-terminus interferes with packing and stability of the complex (43). This can be understood based on their position in the cryo-EM structure: P158 and F160 are both located close to the center of the complex between helix 5 and the beginning of the β-barrel (40). In contrast, the defects introduced by the P132L mutation are much more dramatic, highlighting the importance of this residue in controlling the overall oligomerization state of the protein.

In conclusion, we have now identified the importance of P132 in controlling the oligomerization of CAV1 and the impact of a disease mutation at this site. These findings provide a molecular framework for understanding how defects in caveolins ultimately influence the assembly of caveolar domains and new insights into the fundamental processes that control caveolae biogenesis.

## Materials and Methods

### Cell Culture

*CAV1*^-/-^ MEFs (KO MEFs) were obtained from ATCC and cultured in Dulbecco’s modified Eagle medium (DMEM) containing 10% fetal bovine serum, 1% Pen/Strep at 37° C and 5% CO_2_.

### Constructs and transfections

CAV1-mCherry, CAV1-mycHis and P132L-mycHis for mammalian cell expression and CAV1-His, P132L-His, CAV1-mVenusHis, and P132L-mVenusHis for *E*.*coli* expression were as described previously (38,43,52). The construction of C-terminus HA or myc tagged CAV1 (WT and P132L mutant) expression constructs were based above mentioned CAV1-mycHis or P132L-mycHis plasmids using PCR. The primers used were as follows: CAV1/P132L-HA: CGGGATCCATGTCTGGGGGCAAATACGTAG and CGGAATTCTTAGCTAGCGTAGTCTGGGACGTCGTATGGGTATATTTCTTTCTGCAAGTTGATGCG; CAV1/P132L-myc: CGGGATCCATGTCTGGGGGCAAATACGTAG and CGGAATTCTTACAGATCCTCTTCTGAGATGAGTTTTTGTTCGGGCCCAAGCTTTATTTCTTTCTGCAAGTTGATGCG. Constructs were verified by sequencing.

Transient transfections were performed using Lipofectamine® 2000 (Life Technologies, Carlsbad, CA, USA) as per the manufacturer’s instructions. Cells were transfected 1 day before observation or biochemical analysis.

### Antibodies

Mouse anti-GFP mAb (catalog number 632381) was obtained from Clontech (San Jose, CA, USA). Mouse anti-mCherry mAb (catalog number NBP1-96752) was obtained from NOVUS (Centenial, CO, USA). For Western blots, rabbit anti c-Myc pAb (catalog number sc-789) was obtained from Santa Cruz Biotechnology (Dallas, TX, USA). Mouse anti c-Myc mAb (clone 9B11, #2276) and mouse anti HA mAb (6E2) (catalogue number 2367) were obtained from Cell Signaling Technology (Danvers, MA, USA). Rabbit anti-CAV1 pAb (catalog number 610060), mouse anti-flotillin-1 mAb (catalog number 610820) and mouse anti-calnexin mouse mAb (catalog number 610523) were purchased from BD Transduction Laboratories (Torrey Pines, CA, USA). Rabbit anti-6X His pAb (catalog number 137839), rabbit anti-cavin1 pAb (catalog number 76919) and goat anti-EHD2 pAb (catalog number 23935) were purchased from Abcam (Waltham, MA, USA). Rabbit anti-PACSIN2 pAb (catalog number AP8088b) was purchased from Abgent (San Diego, CA, USA). For Western blotting fluorescently conjugated secondary antibodies and blocking buffer were obtained from LI-COR Biosciences (Lincoln, NE, USA). For immunofluorescence assays, Alexa labeled secondary antibodies were obtained from Life Technologies (Carlsbad, CA, USA).

### Immunofluorescence microscopy

CAV1^-/-^ MEFs were seeded at 150,000 cells/plate in MatTek dishes and transfected with Lipofectamine as per the manufacturer’s instructions. 24 h post transfection, cells were rinsed 2x with PBS and fixed for 12 min in 4% PFA in PBS at room temperature (RT). After another 3x rinsing in PBS containing 100 mM glycine, the cells were blocked and permeabilized with blocking buffer (0.1% saponin in PBS containing 0.5% BSA) at RT for 1h. The cells were stained with anti-Myc antibody (9B11, 1:1000 dilution in blocking buffer) plus antibodies against respective caveolin accessory proteins as indicated in figures (rabbit anti-cavin1, 1:200, Abcam 76919; goat anti-EHD2, 1:200, Abcam 23935; rabbit anti-PACSIN2, 1:200, Abgent AP8088b) at 4°C overnight. Glass-bottom dishes were then rinsed 3x with PBS containing 0.01% Triton X-100 and then incubated in a 1:500 dilution of Alexa 488 and Alexa 647-conjugated secondary antibodies (EHD2 sample: 488 donkey anti mouse IgG, 647 donkey anti goat IgG; cavin1 and PACSIN2 samples: 488 goat anti mouse IgG, 647 goat anti rabbit IgG). After another 3x rinse in PBS containing 0.01% Triton X-100, 1 ml PBS was added to each dish.

For AiryScan imaging, cell samples were imaged using a Zeiss LSM 880 confocal microscope (Jena, Germany) with a Plan-Apochromat 63x/1.4 oil DIC M27 objective, in PBS, at RT. Signals from all three channels were collected with the AiryScan detector in superresolution, frame switching mode. For quantitation, images were taken with the same objective and a 6x zoom factor, in confocal mode. Signals from the far-red channel (Alexa 647) were collected with a PMT detector, while signals from the green (Alexa 488) and red (Alexa 546) channels were collected with a 34-channel GAsP spectral detector. All images were taken in 16 bits format. Contrasts were adjusted linearly with ImageJ. Images were then exported as JPGs.

For total internal reflection fluorescence microscopy, samples were imaged using a Leica Thunder Imager (Wetzlar, Germany) equipped with a TIRF system and the LAX operating software, in 3-channel mode (647/546/488), at RT. A HC PL APO 100x/1.47 oil objective and a Leica-DFC9000GTC-VSC11976 camera (Wetzlar, Germany) were used and the penetration depth was 110 nm.

Colocalization analysis was performed as described before (44) using Macbiophotonics ImageJ with the “Intensity Correlation Analysis” plugin. In brief, images used for colocalization analysis were taken with a 6X zoom factor at 22.5 µm × 22.5 µm dimensions. For each set of comparisons, 90 images from at least three independent experiments were used for analysis. Pearson’s correlation coefficients are reported as the mean ± standard error for all the images. A one-way ANOVA with Tukey’s test (≥3 groups) or an unpaired Student’s t test (2 groups) was used to calculate P-values.

### Electrophoresis

Blue Native-PAGE was performed as descripted before (38) with the NativePAGE™ Bis-Tris Gel System (Life Technologies, Carlsbad, CA, USA). Total membranes of *E. coli* were lysed at 4° C for 30 min with lysis buffer [NativePAGE™ 1X Sample Buffer, complete protease inhibitor cocktail from Roche, and 2% C12M (*n*-dodecyl-β-maltopyranoside) (Anatrace, Maumee, OH, USA)], then spun at 13,100 rpm, 4° C. The supernatant was used for the following analysis. 4-16% NativePAGE™ gels (Life Technologies, Carlsbad, CA, USA) were used for the protein separation. Equal amounts of protein were loaded. The molecular weight was evaluated using NativeMark™ unstained protein standards.

SDS-PAGE electrophoresis was conducted by using Novex® NuPAGE® SDS-PAGE Gel System (Life Technologies, Carlsbad, CA, USA). NuPAGE® 4–12% Bis-Tris gels (Life Technologies, Carlsbad, CA, USA) were used for the protein separation. SeeBlue® Plus2 Pre-stained Protein was used to evaluate the molecular weight.

### Western blotting

The electrophoretic transfer was conducted by using the Mini Trans-Blot® Electrophoretic Transfer system (Bio-Rad, Hercules, CA, USA). For Blue Native-PAGE, the PVDF membranes (Millipore Sigma, Burlington, MA, USA) were de-stained with methanol and washed with TBS (Tris-buffered saline) buffer. An LI-COR Odyssey infrared imaging system (LI-COR Biosciences, Lincoln, NE, USA) was used for the signal detecting of blots. Quantification of Western blot images was performed using IMAGEJ.

### Caveolin Complexes Fractionation

Caveolin complexes were fractionated by velocity gradient centrifugation as described before (20,38). For each fractionation, 4 × 10^6^ *Cav1*^-/-^ MEFs were lysed for 20 min in 330 μL 0.5% TX-100 (or 0.4% SDS plus 0.2% TX-100) in TNE [100 mM NaCl, 20 m M Tris–HCl pH 7.5, 5 mM ethylenediaminetetraacetic acid (EDTA)] buffer, supplemented with ‘Complete’ protease inhibitors cocktail (Roche, Basel, Switzerland) at RT. PNSs (Postnuclear supernatants) were prepared by conducting a 5-min centrifugation at 1100 × g. 300μL of the PNS was recovered and loaded onto linear 10–40% linear sucrose gradients. The sucrose solution was prepared with TNE buffer with protease inhibitors cocktail (Roche, Basel, Switzerland). Sucrose gradients were centrifuged in an SW55 rotor using a Optima™ LE-80K Ultracentrifuge (Beckman Coulter, Pasadena, CA, USA)) for 5 hours at 48,000 rpm and 4° C. 14 equal volume (about 360μL) fractions were harvested from the top and analyzed by SDS-PAGE/Western Blot with 10μL loading from each fraction.

### DRM fractionation

Caveolae-enriched DRMs were fractionated as described (38). Approximemtly1.6 × 10^7^ *Cav1*^-/-^ MEFs were suspended in 300 μl precooled 0.5% TX100 in TNE buffer, supplemented with ‘Complete’ protease inhibitors cocktail (Roche, Basel, Switzerland). Cell suspensions were homogenized by passing 10 times through a precooled 1-ml syringe with a 27-gauge stainless steel needle (BD Biosciences, Franklin Lakes, NJ). The homogenate was adjusted to about 40% sucrose by the addition of 700μl of 60% sucrose prepared in TNE and placed at the bottom of an ultracentrifuge tube. A 5 to 30% linear sucrose gradient was formed above the homogenate and centrifuged at 40,100 rpm and 4° C for 16 hours in a SW55 rotor using an OptimaTM LE-80K Ultracentrifuge (Beckman Coulter, Pasadena, CA). Fourteen 360 μl fractions were collected from the top and analyzed by SDS–PAGE/Western blot with an equal loading volume. Western blots were images and quantified as indicated above.

### Expression of caveolin in *E. coli* and purification of CAV1 complexes

Protein expression and purification was conducted as described with minor modifications (43). In brief, caveolin proteins were expressed in *E. coli* BL21 using the auto-induction expression system (67). First, MDG starter of monoclonal bacteria was cultured at 37°C and 250 rpm for 20 hours, then auto-inducing ZYM-5052 media was used to enlarge culture at 25°C and 300 rpm for 24 hours. *E. coli* cells were washed with 0.9% NaCl and then were resuspended with buffer (200 mM NaCl and 20 mM Tris-HCl, pH 8.0).

Bacterial cells were homogenized with a French press pressure homogenizer, and 1 mM PMSF and DTT were added just prior to homogenization. A 15 min centrifugation at 9000 rpm and 4°C was conducted to remove large cell debris, then total membranes were pelleted at 40,000 rpm (Ti-45 rotor, Beckman Coulter, Pasadena, CA, USA) and 4°C for 1 hour. Membrane pellets were homogenized with Dounce tissue grinder in a buffer composed of 200 mM NaCl, 20 mM Tris-HCl (pH 8.0), and 1 mM DTT. To solubilize caveolin proteins from membranes, 10% C12M (Anatrace, Maumee, OH, USA) stock solution was mixed into membrane homogenate to a final concentration of 2%, and the mixture was slowly stirred for 2 hours at 4°C. Insoluble material was pelleted down by centrifugation at 42,000 rpm (Ti-50.2 rotor) for 35 min, and the supernatant was used for nickel Sepharose–based affinity purification. The caveolin-containing eluate was concentrated and further separated by size exclusion chromatography using a Superpose 6 Increase 10/300 GL column (GE Healthcare, Boston, MA, USA) in buffer containing 200 mM NaCl, 20 mM Tris-HCl (pH 8.0), 1 mM DTT, and 0.05% C12M.

### Negative stain electron microscopy and data processing

Negative stain EM was performed using established methods (68). In brief, 200-mesh copper grids covered with carbon-coated collodion film (EMS, Hatfield, PA, USA) were glow discharged for 30 s at 10 mA in a PELCO easiGlow™ glow discharge unit (Fresno, CA, USA). Aliquots (3.5 μl) of purified sample were adsorbed to the grids and incubated for 1 minute at room temperature. Samples were then washed with 2 drops of water and stained with 2 successive drops of 0.7% (w/v) uranyl formate (EMS, Hatfield, PA, USA) followed by blotting until dry. Samples were visualized on a Morgagni transmission electron microscope equipped with a field emission gun operating at an accelerating voltage of 100 keV (Thermo Fisher Scientific, Waltham, MA, USA) at a nominal magnification of 22,000x (2.1 Å per pixel).

The negative stain datasets were collected using a Tecnai Spirit T12 transmission electron microscope operated at 120keV (Thermo Fisher Scientific, Waltham, MA, USA). Datasets were collected at a nominal magnification of ×26,000 (2.34 Å per pixel) except for the P132L-mVenus and P132L-mVenus + CAV1-His datasets, which were collected at a nominal magnification of x42,000 (1.45Å per pixel). Sample data was collected using Leginon software on a 4 k × 4 k Rio complementary metal-oxide semiconductor camera (Gatan, Pleasanton, CA) at −1.5-μm defocus value (69). Images were manually curated. All data processing was carried out in Relion 3.1.0 (70). About 1,000 particles were picked manually and 2D classified. Clear resulting classes were selected and used as references for particle selection on all images. Particles were extracted with a 128 pixel (2.34 Å per pixel datasets) or 208 pixel (1.45Å per pixel datasets) box size (30nm by 30nm boxes). The extracted particles were then 2D classified. The P132L-His and P132L-mVenus datasets consisted of 35,405 and 12,634 particles, respectively. The P132L-mVenus + P132L-His, P132L-His + CAV1-mVenus, and P132L-mVenus + CAV1-His datasets had 47,711, 51,339, and 12,015 particles.The CAV1-His, CAV1-mVenus, and CAV1-mVenus + CAV1-His datasets had 31,956, 9,370, and 42,197 particles.

### Symmetric ddG calculations

The experimental CAV1 structure was used as the starting point for all the calculations (PDB 7SC0). Membrane coordinates of this structure were calculated with the PPM (Positioning of Proteins in Membranes) server (71). A single chain from this structure was used as the input for the symmetric *FastRelax* calculations in Rosetta 3.10. 11-fold symmetry files were created with Rosetta 3.10 based on the experimental structure, and this symmetry was imposed on the monomeric unit for all the following calculations. The position 132 was mutated into all 20 amino acids (including P132P) to screen for the effect of different mutations at this site. Specifically, the amino acid at this site was first mutated using the *MutateResidue* mover of Rosetta 3.10, followed by a *FastRelax* calculation to minimize the system with restraints on the system (0.5 Å deviation was allowed). All the calculations were run with the membrane score function *mpframework_smooth_fa_2012*.

### Asymmetric ddG calculations

The 11-meric experimental CAV1 structure was used for the calculations (PDB 7SC0). In the first step, all 20 amino acid mutations were introduced at the position 132, and the corresponding Rosetta energy difference between these mutations and the P132P self-mutation was measured to assess their effects. All calculations were run with the *mp2012* membrane score function with the cav-1 pre-aligned in the membrane through the PPM server (71). Only backbone motions and side chain repacking were allowed for the residues within 8 Å of position 132.

## This article contains supporting information

## Acknowledgements

The University of Michigan Cryo-EM Facility (U-M Cryo-EM) has received generous support from the U-M Life Sciences Institute and the U-M Biosciences Initiative.

## Author contributions

Conceptualization: BH, JM, MDO, AKK

Methodology: BH, AG

Investigation: BH, AG, SC, TW, JP, AT, AD, LC, YP

Data curation: SC, JP

Formal analysis: AG, SC, TW, JP

Writing-original draft: BH, AG, SC, TW, EK, MDO, AKK

Writing-review and editing: BH, AG, SC, TW, JP, AT, AD, LC, YP, HSM, EK, MDO, JM, AKK

Validation: BH, AT, AD, TW

Visualization: BH, AG, SC, TW, JP, EK

Supervision: AKK, MDO, EK, JM

Project administration: AKK, MDO, JM

Funding acquisition: AKK, MDO, JM

## Funding and additional information

This work was supported by National Institutes of Health grant R01 HL144131 (AKK and MDO), NIH R01 HL111259 (AKK), NIH R01 GM106720 (AKK), National Institutes of Health grant S10OD020011 (MDO), National Institutes of Health grant S10OD030275 (MDO), National Institutes of Health grant T-32-GM007315 (SC), American Heart Association grant 90705 (SC), National Institutes of Health grant R01GM080403 (JM), National Institutes of Health grant R01HL122010 (JM), National Institutes of Health grant R01GM129261 (JM), and Humboldt Professorship of the Alexander von Humboldt Foundation (JM). The content is solely the responsibility of the authors and does not necessarily represent the official views of the National Institutes of Health.

## References

1. Parton, R. G., and del Pozo, M. A. (2013) Caveolae as plasma membrane sensors, protectors and organizers. Nat Rev Mol Cell Biol 14, 98–112

2. Ariotti, N., and Parton, R. G. (2013) SnapShot: caveolae, caveolins, and cavins. Cell 154, 704–704 e701

3. Hansen, C. G., and Nichols, B. J. (2010) Exploring the caves: cavins, caveolins and caveolae. Trends Cell Biol 20, 177–186

4. Razani, B., and Lisanti, M. P. (2001) Caveolin-deficient mice: insights into caveolar function human disease. J Clin Invest 108, 1553–1561

5. Walser, P. J., Ariotti, N., Howes, M., Ferguson, C., Webb, R., Schwudke, D., Leneva, N., Cho, K. J., Cooper, L., Rae, J., Floetenmeyer, M., Oorschot, V. M., Skoglund, U., Simons, K., Hancock, J. F., and Parton, R. G. (2012) Constitutive formation of caveolae in a bacterium. Cell 150, 752–763

6. Ariotti, N., Rae, J., Leneva, N., Ferguson, C., Loo, D., Okano, S., Hill, M. M., Walser, P., Collins, B. M., and Parton, R. G. (2015) Molecular characterization of caveolin-induced membrane curvature. J Biol Chem 290, 24875–24890

7. Lamaze, C., Tardif, N., Dewulf, M., Vassilopoulos, S., and Blouin, C. M. (2017) The caveolae dress code: structure and signaling. Curr Opin Cell Biol 47, 117–125

8. Parton, R. G., McMahon, K. A., and Wu, Y. (2020) Caveolae: formation, dynamics, and function. Curr Opin Cell Biol 65, 8–16

9. Andrade, V., Bai, J., Gupta-Rossi, N., Jimenez, A. J., Delevoye, C., Lamaze, C., and Echard, A. (2022) Caveolae promote successful abscission by controlling intercellular bridge tension during cytokinesis. Sci Adv 8, eabm5095

10. Parton, R. G. (2018) Caveolae: structure, function, and relationship to disease. Annu Rev Cell Dev Biol 34, 111–136

11. Plucinsky, S., and Glover, K. (2017) The C-terminal domain of caveolin-1 and pulmonary arterial hypertension: An emerging relationship. J Rare Dis Res Treat. 2, 44–48

12. Patni, N., and Garg, A. (2015) Congenital generalized lipodystrophies-new insights into metabolic dysfunction. Nat Rev Endocrinol 11, 522–534

13. Williams, T. M., and Lisanti, M. P. (2004) The Caveolin genes: from cell biology to medicine. Ann Med 36, 584–595

14. Lamaze, C., and Torrino, S. (2015) Caveolae and cancer: A new mechanical perspective. Biomed J 38, 367–379

15. Yin, H., Liu, T., Zhang, Y., and Yang, B. (2016) Caveolin proteins: a molecular insight into disease. Front Med 10, 397–404

16. Mathew, R. (2021) Critical Role of Caveolin-1 Loss/Dysfunction in Pulmonary Hypertension. Med Sci (Basel) 9

17. Monier, S., Parton, R. G., Vogel, F., Behlke, J., Henske, A., and Kurzchalia, T. V. (1995) VIP21-caveolin, a membrane protein constituent of the caveolar coat, oligomerizes in vivo and in vitro. Mol. Biol. Cell 6, 911–927

18. Sargiacomo, M., Scherer, P. E., Tang, Z., Kubler, E., Song, K. S., Sanders, M. C., and Lisanti, M. P. (1995) Oligomeric structure of caveolin: implications for caveolae membrane organization. Proc Natl Acad Sci U S A 92, 9407–9411

19. Ren, X., Ostermeyer, A. G., Ramcharan, L. T., Zeng, Y., Lublin, D. M., and Brown, D. A. (2004) Conformational defects slow Golgi exit, block oligomerization, and reduce raft affinity of caveolin-1 mutant proteins. Mol Biol Cell 15, 4556–4567

20. Hayer, A., Stoeber, M., Bissig, C., and Helenius, A. (2010) Biogenesis of caveolae: stepwise assembly of large caveolin and cavin complexes. Traffic 11, 361–382

21. Han, B., Copeland, C. A., Tiwari, A., and Kenworthy, A. K. (2016) Assembly and turnover of caveolae: what do we really know? Front Cell Dev Biol 4, 68

22. Parton, R. G., Tillu, V., McMahon, K. A., and Collins, B. M. (2021) Key phases in the formation of caveolae. Curr Opin Cell Biol 71, 7–14

23. Busija, A. R., Patel, H. H., and Insel, P. A. (2017) Caveolins and cavins in the trafficking, maturation, and degradation of caveolae: implications for cell physiology. Am J Physiol Cell Physiol 312, C459–C477

24. Matthaeus, C., and Taraska, J. W. (2020) Energy and Dynamics of Caveolae Trafficking. Front Cell Dev Biol 8, 614472

25. Tang, Z., Okamoto, T., Boontrakulpoontawee, P., Katada, T., Otsuka, A. J., and Lisanti, M. P. (1997) Identification, sequence, and expression of an invertebrate caveolin gene family from the nematode Caenorhabditis elegans. Implications for the molecular evolution of mammalian caveolin genes. J Biol Chem 272, 2437–2445

26. Kirkham, M., Nixon, S. J., Howes, M. T., Abi-Rached, L., Wakeham, D. E., Hanzal-Bayer, M., Ferguson, C., Hill, M. M., Fernandez-Rojo, M., Brown, D. A., Hancock, J. F., Brodsky, F. M., and Parton, R. G. (2008) Evolutionary analysis and molecular dissection of caveola biogenesis. J Cell Sci 121, 2075–2086

27. Hayashi, K., Matsuda, S., Machida, K., Yamamoto, T., Fukuda, Y., Nimura, Y., Hayakawa, T., and Hamaguchi, M. (2001) Invasion activating caveolin-1 mutation in human scirrhous breast cancers. Cancer Res 61, 2361–2364

28. Shatz, M., Lustig, G., Reich, R., and Liscovitch, M. (2010) Caveolin-1 mutants P132L and Y14F are dominant negative regulators of invasion, migration and aggregation in H1299 lung cancer cells. Exp Cell Res 316, 1748–1762

29. Bonuccelli, G., Casimiro, M. C., Sotgia, F., Wang, C., Liu, M., Katiyar, S., Zhou, J., Dew, E., Capozza, F., Daumer, K. M., Minetti, C., Milliman, J. N., Alpy, F., Rio, M. C., Tomasetto, C., Mercier, I., Flomenberg, N., Frank, P. G., Pestell, R. G., and Lisanti, M. P. (2009) Caveolin-1 (P132L), a common breast cancer mutation, confers mammary cell invasiveness and defines a novel stem cell/metastasis-associated gene signature. Am J Pathol 174, 1650–1662

30. Koike, S., Kodera, Y., Nakao, A., Iwata, H., and Yatabe, Y. (2010) Absence of the caveolin-1 P132L mutation in cancers of the breast and other organs. J Mol Diagn 12, 712–717

31. Lacroix-Triki, M., Geyer, F. C., and Reis-Filho, J. S. (2010) Caveolin-1 P132L mutation in human cancers: 1 CAVeat to be voiced. J Mol Diagn 12, 562–565

32. Ferraldeschi, R., Latif, A., Clarke, R. B., Spence, K., Ashton, G., O’Sullivan, J., Evans, D. G., Howell, A., and Newman, W. G. (2012) Lack of caveolin-1 (P132L) somatic mutations in breast cancer. Breast Cancer Res Treat 132, 1185–1186

33. Patani, N., Lambros, M. B., Natrajan, R., Dedes, K. J., Geyer, F. C., Ward, E., Martin, L. A., Dowsett, M., and Reis-Filho, J. S. (2012) Non-existence of caveolin-1 gene mutations in human breast cancer. Breast Cancer Res Treat 131, 307–310

34. Minetti, C., Sotgia, F., Bruno, C., Scartezzini, P., Broda, P., Bado, M., Masetti, E., Mazzocco, M., Egeo, A., Donati, M. A., Volonte, D., Galbiati, F., Cordone, G., Bricarelli, F. D., Lisanti, M. P., and Zara, F. (1998) Mutations in the caveolin-3 gene cause autosomal dominant limb-girdle muscular dystrophy. Nat Genet 18, 365–368

35. Galbiati, F., Volonte, D., Minetti, C., Chu, J. B., and Lisanti, M. P. (1999) Phenotypic behavior of caveolin-3 mutations that cause autosomal dominant limb girdle muscular dystrophy (LGMD-1C). Retention of LGMD-1C caveolin-3 mutants within the golgi complex. J Biol Chem 274, 25632–25641

36. Carbone, I., Bruno, C., Sotgia, F., Bado, M., Broda, P., Masetti, E., Panella, A., Zara, F., Bricarelli, F. D., Cordone, G., Lisanti, M. P., and Minetti, C. (2000) Mutation in the CAV3 gene causes partial caveolin-3 deficiency and hyperCKemia. Neurology 54, 1373–1376

37. Lee, H., Park, D. S., Razani, B., Russell, R. G., Pestell, R. G., and Lisanti, M. P. (2002) Caveolin-1 mutations (P132L and null) and the pathogenesis of breast cancer: caveolin-1 (P132L) behaves in a dominant-negative manner and caveolin-1 (-/-) null mice show mammary epithelial cell hyperplasia. Am J Pathol 161, 1357–1369

38. Han, B., Tiwari, A., and Kenworthy, A. K. (2015) Tagging strategies strongly affect the fate of overexpressed caveolin-1. Traffic 16, 417–438

39. Rieth, M. D., Lee, J., and Glover, K. J. (2012) Probing the caveolin-1 P132L mutant: critical insights into its oligomeric behavior and structure. Biochemistry

40. Porta, J. C., Han, B., Gulsevin, A., Chung, J. M., Peskova, Y., Connolly, S., McHaourab, H. S., Meiler, J., Karakas, E., Kenworthy, A. K., and Ohi, M. D. (2022) Molecular architecture of the human caveolin-1 complex. Sci Adv 8, eabn7232

41. Scherer, P. E., Okamoto, T., Chun, M., Nishimoto, I., Lodish, H. F., and Lisanti, M. P. (1996) Identification, sequence, and expression of caveolin-2 defines a caveolin gene family. Proc Natl Acad Sci U S A 93, 131–135

42. Tang, Z., Scherer, P. E., Okamoto, T., Song, K., Chu, C., Kohtz, D. S., Nishimoto, I., Lodish, H. F., and Lisanti, M. P. (1996) Molecular cloning of caveolin-3, a novel member of the caveolin gene family expressed predominantly in muscle. J Biol Chem 271, 2255–2261

43. Han, B., Porta, J. C., Hanks, J. L., Peskova, Y., Binshtein, E., Dryden, K., Claxton, D. P., McHaourab, H. S., Karakas, E., Ohi, M. D., and Kenworthy, A. K. (2020) Structure and assembly of CAV1 8S complexes revealed by single particle electron microscopy. Sci Adv 6, eabc6185

44. Han, B., Copeland, C. A., Kawano, Y., Rosenzweig, E. B., Austin, E. D., Shahmirzadi, L., Tang, S., Raghunathan, K., Chung, W. K., and Kenworthy, A. K. (2016) Characterization of a caveolin-1 mutation associated with both pulmonary arterial hypertension and congenital generalized lipodystrophy. Traffic 17, 1297–1312

45. Copeland, C. A., Han, B., Tiwari, A., Austin, E. D., Loyd, J. E., West, J. D., and Kenworthy, A. K. (2017) A disease-associated frameshift mutation in caveolin-1 disrupts caveolae formation and function through introduction of a de novo ER retention signal. Mol Biol Cell 28, 3095–3111

46. Song, K. S., Tang, Z. L., Li, S. W., and Lisanti, M. P. (1997) Mutational analysis of the properties of caveolin-1. A novel role for the C-terminal domain in mediating homo-typic caveolin-caveolin interactions. J. Biol. Chem. 272, 4398–4403

47. Senju, Y., Itoh, Y., Takano, K., Hamada, S., and Suetsugu, S. (2011) Essential role of PACSIN2/syndapin-II in caveolae membrane sculpting. J Cell Sci 124, 2032–2040

48. Moren, B., Shah, C., Howes, M. T., Schieber, N. L., McMahon, H. T., Parton, R. G., Daumke, O., and Lundmark, R. (2012) EHD2 regulates caveolar dynamics via ATP-driven targeting and oligomerization. Mol Biol Cell 23, 1316–1329

49. Hubert, M., Larsson, E., and Lundmark, R. (2020) Keeping in touch with the membrane; protein-and lipid-mediated confinement of caveolae to the cell surface. Biochem Soc Trans 48, 155–163

50. Ludwig, A., Howard, G., Mendoza-Topaz, C., Deerinck, T., Mackey, M., Sandin, S., Ellisman, M. H., and Nichols, B. J. (2013) Molecular composition and ultrastructure of the caveolar coat complex. PLoS Biol 11, e1001640

51. Matthaeus, C., Sochacki, K. A., Dickey, A., Puchkov, D., Haucke, V., Lehmann, M., and Taraska, J. W. (2022) The molecular organization of flat and curved caveolae indicates bendable structural units at the plasma membrane. bioRxiv, 2022.2003.2031.486578

52. Hanson, C. A., Drake, K. R., Baird, M. A., Han, B., Kraft, L. J., Davidson, M. W., and Kenworthy, A. K. (2013) Overexpression of caveolin-1 is sufficient to phenocopy the behavior of a disease-associated mutant. Traffic 14, 663–677

53. Tiwari, A., Copeland, C. A., Han, B., Hanson, C. A., Raghunathan, K., and Kenworthy, A. K. (2016) Caveolin-1 is an aggresome-inducing protein. Sci Rep 6, 38681

54. Pradhan, B. S., and Proszynski, T. J. (2020) A role for caveolin-3 in the pathogenesis of muscular dystrophies. Int J Mol Sci 21

55. Mercier, I., Jasmin, J. F., Pavlides, S., Minetti, C., Flomenberg, N., Pestell, R. G., Frank, P. G., Sotgia, F., and Lisanti, M. P. (2009) Clinical and translational implications of the caveolin gene family: lessons from mouse models and human genetic disorders. Lab Invest 89, 614–623

56. Shah, D. S., Nisr, R. B., Stretton, C., Krasteva-Christ, G., and Hundal, H. S. (2020) Caveolin-3 deficiency associated with the dystrophy P104L mutation impairs skeletal muscle mitochondrial form and function. J Cachexia Sarcopenia Muscle 11, 838–858

57. Woodman, S. E., Sotgia, F., Galbiati, F., Minetti, C., and Lisanti, M. P. (2004) Caveolinopathies: mutations in caveolin-3 cause four distinct autosomal dominant muscle diseases. Neurology 62, 538–543

58. Garg, A., Kircher, M., Del Campo, M., Amato, R. S., and Agarwal, A. K. (2015) Whole exome sequencing identifies de novo heterozygous CAV1 mutations associated with a novel neonatal onset lipodystrophy syndrome. Am J Med Genet A 167, 1796–1806

59. Schrauwen, I., Szelinger, S., Siniard, A. L., Kurdoglu, A., Corneveaux, J. J., Malenica, I., Richholt, R., Van Camp, G., De Both, M., Swaminathan, S., Turk, M., Ramsey, K., Craig, D. W., Narayanan, V., and Huentelman, M. J. (2015) A frame-shift mutation in CAV1 is associated with a severe neonatal progeroid and lipodystrophy syndrome. PLoS One 10, e0131797

60. Marsboom, G., Chen, Z., Yuan, Y., Zhang, Y., Tiruppathi, C., Loyd, J. E., Austin, E. D., Machado, R. F., Minshall, R. D., Rehman, J., and Malik, A. B. (2017) Aberrant caveolin-1-mediated Smad signaling and proliferation identified by analysis of adenine 474 deletion mutation (c.474delA) in patient fibroblasts: a new perspective on the mechanism of pulmonary hypertension. Mol Biol Cell 28, 1177–1185

61. Kim, C. A., Delepine, M., Boutet, E., El Mourabit, H., Le Lay, S., Meier, M., Nemani, M., Bridel, E., Leite, C. C., Bertola, D. R., Semple, R. K., O’Rahilly, S., Dugail, I., Capeau, J., Lathrop, M., and Magre, J. (2008) Association of a homozygous nonsense caveolin-1 mutation with Berardinelli-Seip congenital lipodystrophy. J Clin Endocrinol Metab 93, 1129–1134

62. Cao, H., Alston, L., Ruschman, J., and Hegele, R. A. (2008) Heterozygous CAV1 frameshift mutations (MIM 601047) in patients with atypical partial lipodystrophy and hypertriglyceridemia. Lipids Health Dis 7, 3

63. Karhan, A. N., Zammouri, J., Auclair, M., Capel, E., Apaydin, F. D., Ates, F., Verpont, M. C., Magre, J., Feve, B., Lascols, O., Usta, Y., Jeru, I., and Vigouroux, C. (2021) Biallelic CAV1 null variants induce congenital generalized lipodystrophy with achalasia. Eur J Endocrinol 185, 841–854

64. Sotgia, F., Woodman, S. E., Bonuccelli, G., Capozza, F., Minetti, C., Scherer, P. E., and Lisanti, M. P. (2003) Phenotypic behavior of caveolin-3 R26Q, a mutant associated with hyperCKemia, distal myopathy, and rippling muscle disease. Am J Physiol Cell Physiol 285, C1150–1160

65. Morales-Paytuvi, F., Ruiz-Mirapeix, C., Fajardo, A., Rae, J., Bosch, M., Enrich, C., Collins, B., Parton, R. G., and Pol, A. (2022) Proteostatic regulation of caveolins avoids premature oligomerisation and preserves ER homeostasis. bioRxiv

66. Austin, E. D., Ma, L., LeDuc, C., Berman Rosenzweig, E., Borczuk, A., Phillips, J. A., 3rd, Palomero, T., Sumazin, P., Kim, H. R., Talati, M. H., West, J., Loyd, J. E., and Chung, W. K. (2012) Whole exome sequencing to identify a novel gene (caveolin-1) associated with human pulmonary arterial hypertension. Circ Cardiovasc Genet 5, 336–343

67. Studier, F. W. (2005) Protein production by auto-induction in high density shaking cultures. Protein Expr Purif 41, 207–234

68. Ohi, M., Li, Y., Cheng, Y., and Walz, T. (2004) Negative staining and image classification - powerful tools in modern electron microscopy. Biol Proced Online 6, 23–34

69. Suloway, C., Pulokas, J., Fellmann, D., Cheng, A., Guerra, F., Quispe, J., Stagg, S., Potter, C. S., and Carragher, B. (2005) Automated molecular microscopy: the new Leginon system. J Struct Biol 151, 41–60

70. Zivanov, J., Nakane, T., and Scheres, S. H. W. (2020) Estimation of high-order aberrations and anisotropic magnification from cryo-EM data sets in RELION-3.1. IUCrJ 7, 253–267

71. Lomize, M. A., Pogozheva, I. D., Joo, H., Mosberg, H. I., and Lomize, A. L. (2012) OPM database and PPM web server: resources for positioning of proteins in membranes. Nucleic Acids Res 40, D370–376

