## Supplementary Material for "Structural characterization of a breast cancer-associated mutation in caveolin-1"

**Running title: Structural basis for the P132L mutation in caveolin-1**

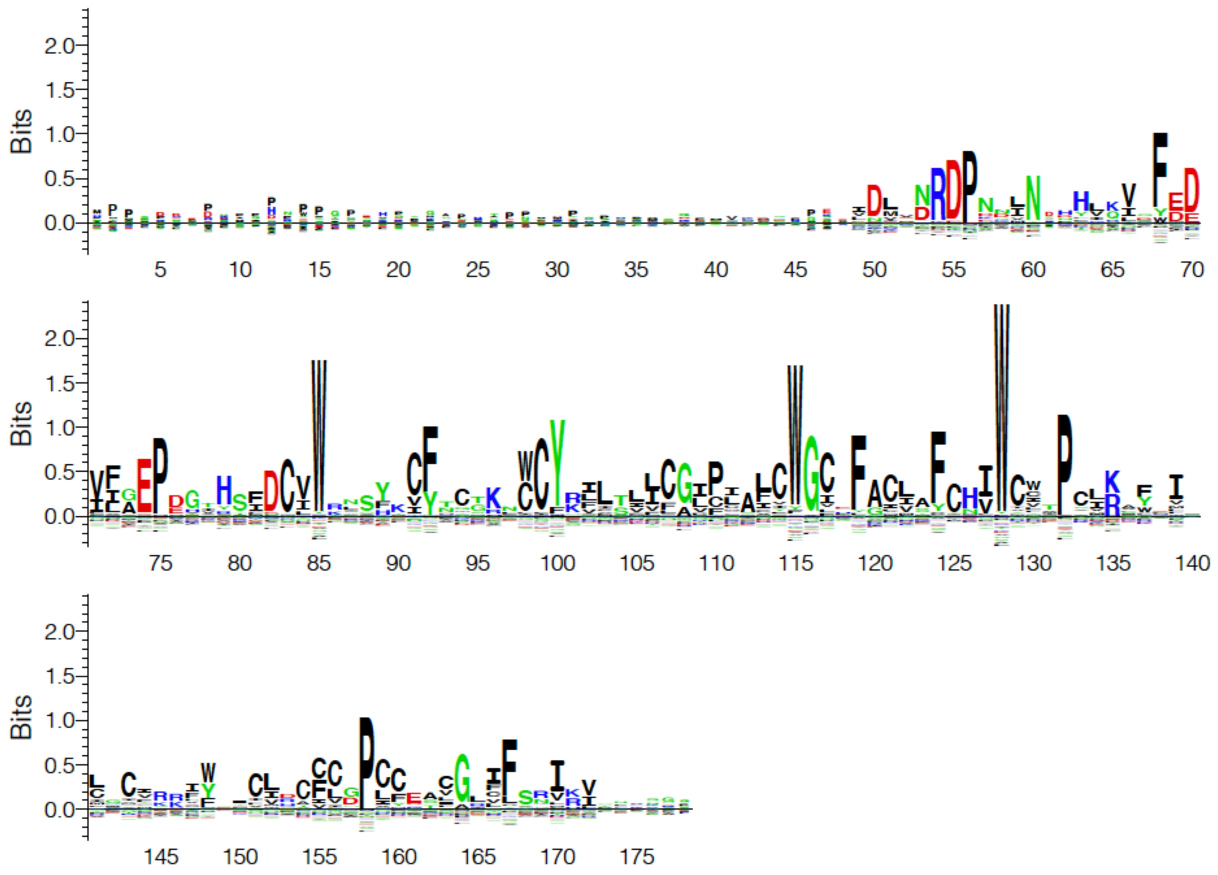


**Figure S1. Sequence logo of CAV1 among metazoan caveolins.** Logo was created with Seq2Logo.

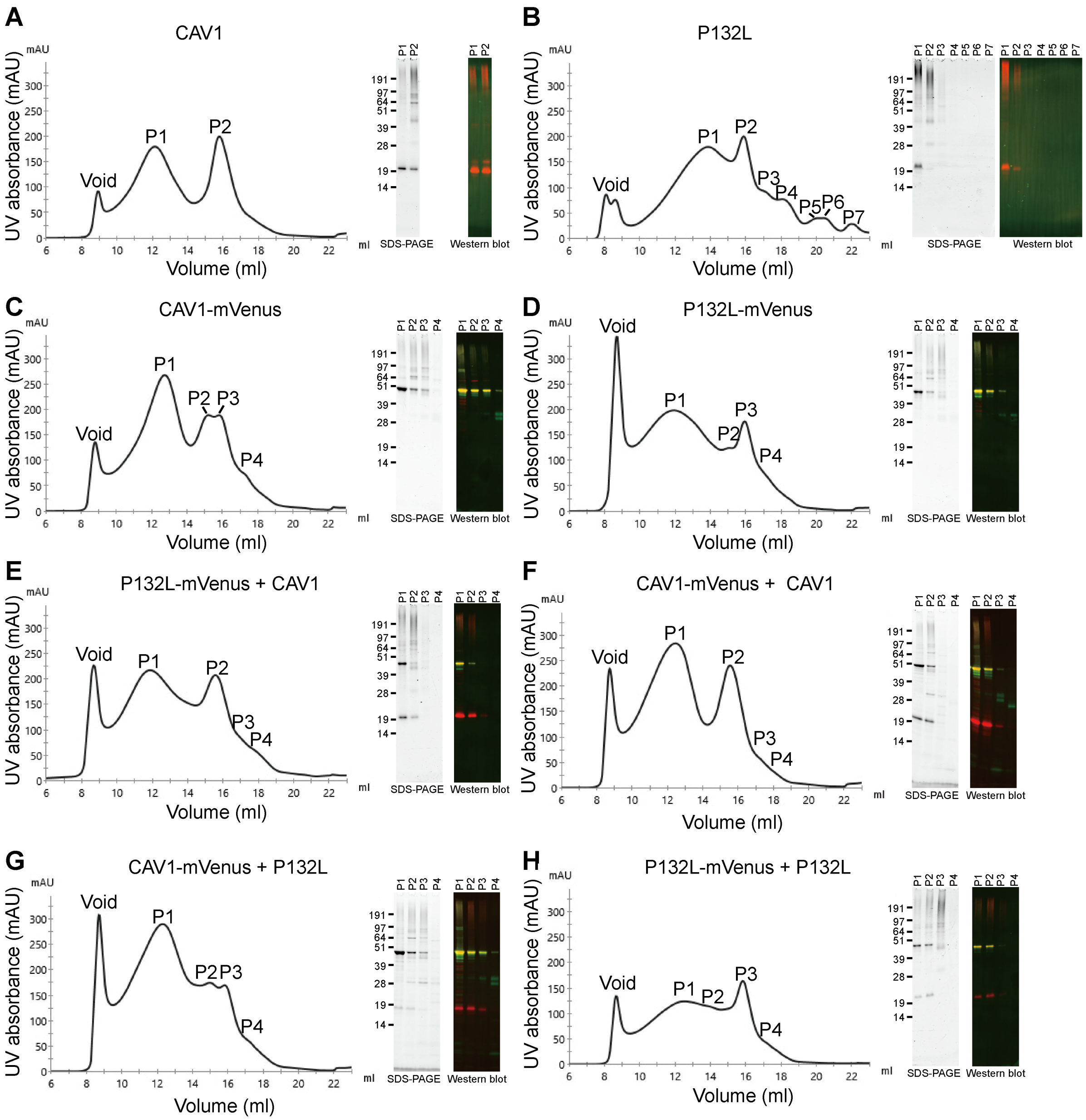


**Figure S2: FPLC traces, SDS-PAGE electrophoresis and Western blotting of CAV1 constructs used in this study.** The indicated Cav1 proteins were purified from *E. coli* membranes and applied to a Superose®6 10/300 GL column. Elution profiles, SDS-PAGE and Western blotting results are shown for (**A**) CAV1, (**B**) P132L, (**C**) CAV1-mVenus, (**D**) P132L-mVenus, (**E**) co-expressed P132L-mVenus and CAV1, (**F**) co-expressed CAV1-mVenus and CAV1, (**G**) co-expressed CAV1-mVenus and P132L, and (**H**) co-expressed P132L and P132L-mVenus. In the Western blotting results, mVenus signal is shown in green and CAV1 signal is in red.

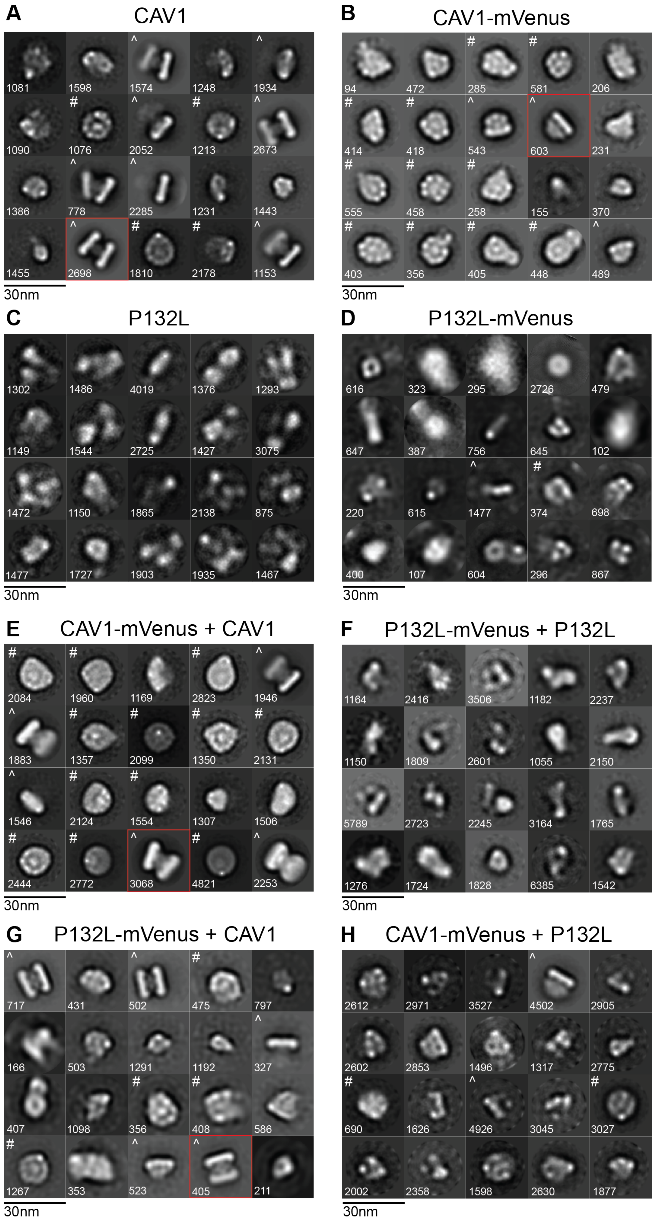


**Figure S3: Negative stain averages of Cav1 constructs.** Negative stain class averages of (**A**) CAV1, (**B**) CAV1-mVenus, (**C**) P132L, (**D**) P132L-mVenus, (**E**) co-expressed CAV1-Venus and CAV1 (**F**), co-expressed P132L-mVenus and P132L, (**G**) co-expressed P132L-mVenus and CAV1, and (**H**) co-expressed CAV1-Venus and P132L. # denotes en face averages, ^ denotes side view averages, and the number of particles per average are noted in the bottom left. Scale bars, 30 nm. Classes in red boxes are displayed in **Figure 4**.

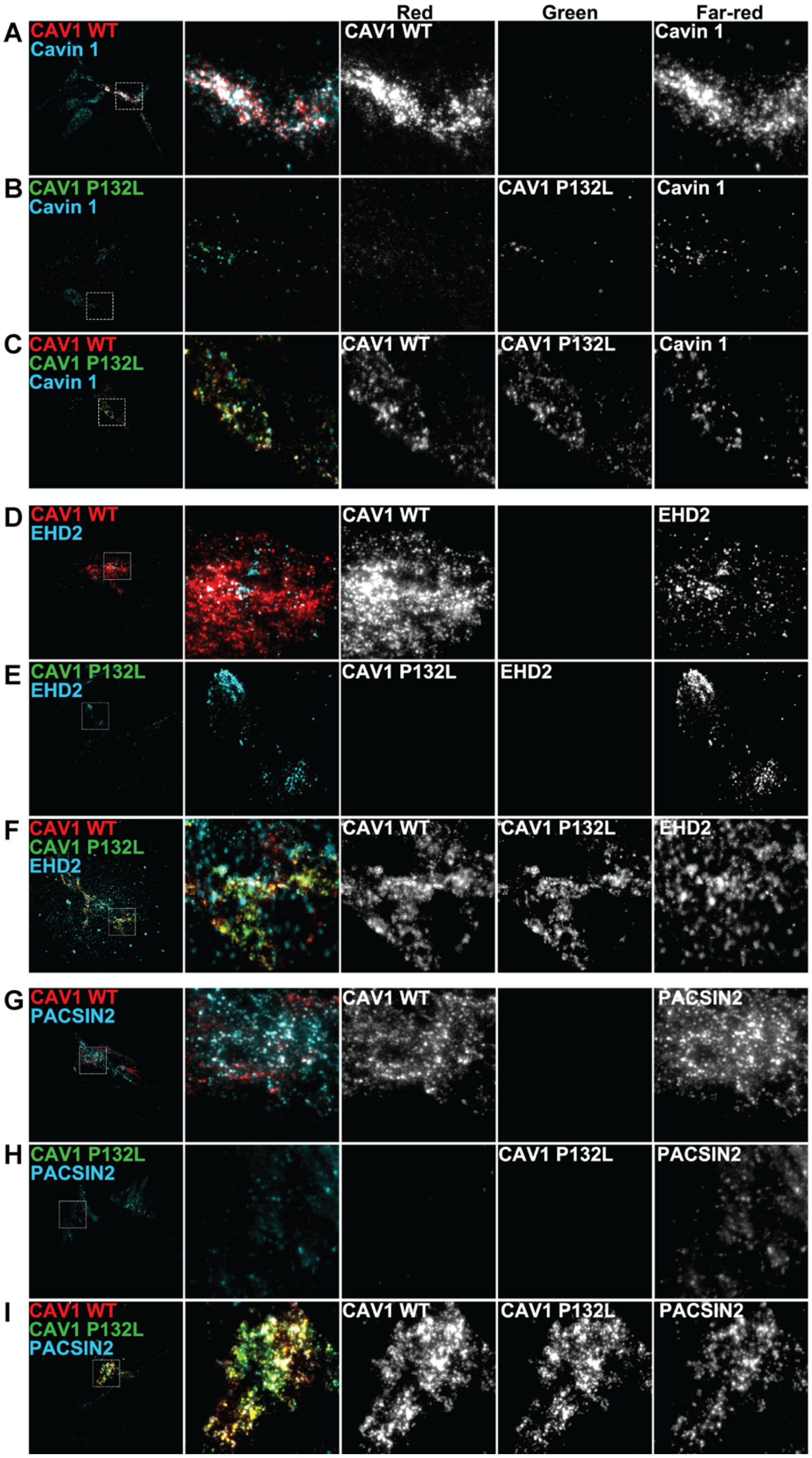


**Figure S4. When co-expressed with WT CAV1, P132L is recruited to cell surface puncta that partially co-localize with cavin-1, EHD2, and PACSIN-2 as visualized using TIRF microscopy.**

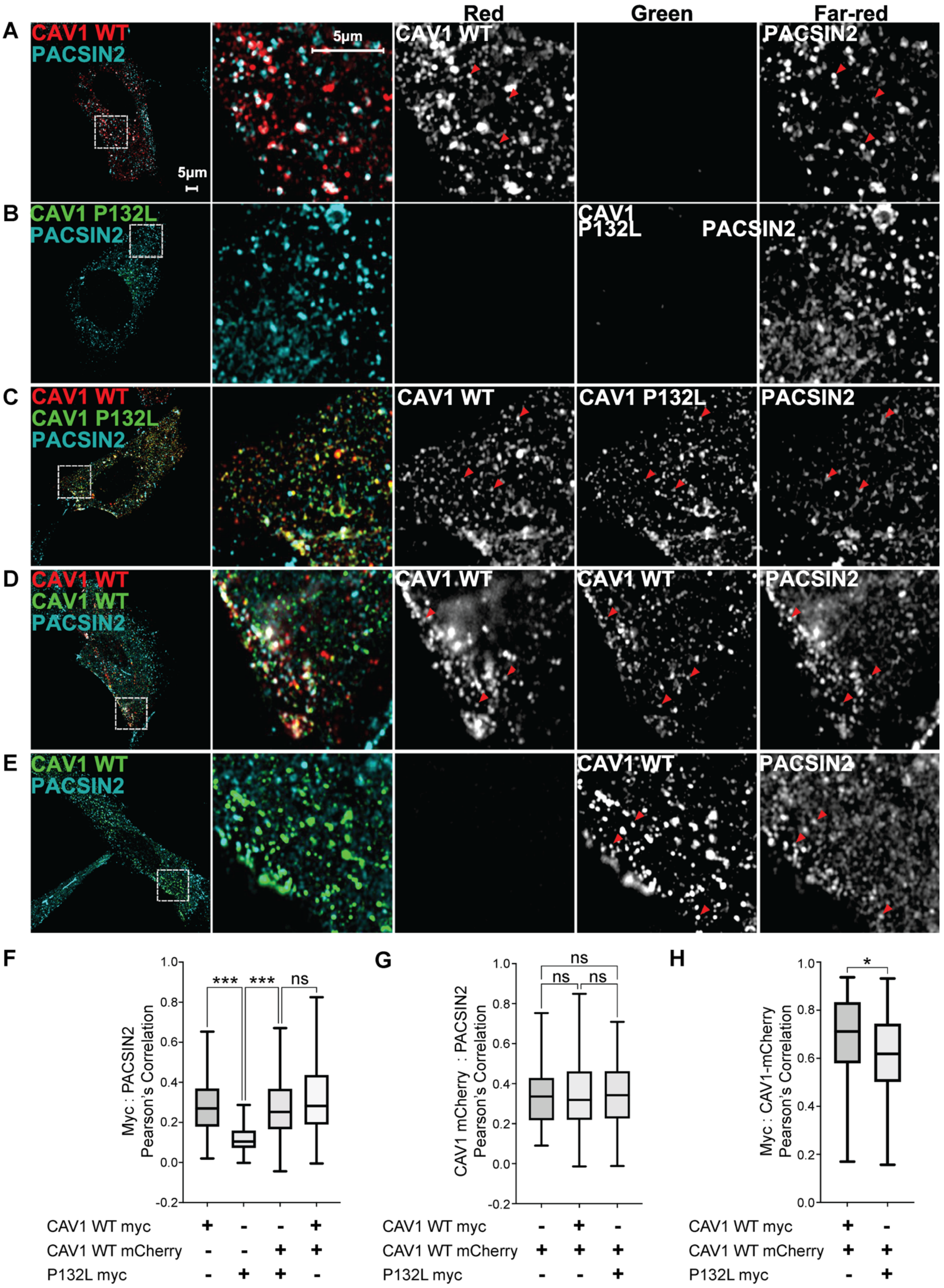

Red
Green
Far-red
A
CAV1 WT
PACSIN2
5μm
5μm
CAV1 WT
PACSIN2
B
CAV1 P132L
PACSIN2
CAV1 P132L
PACSIN2
C
CAV1 WT
CAV1 P132L
PACSIN2
CAV1 WT
CAV1 P132L
PACSIN2
D
CAV1 WT
CAV1 WT
PACSIN2
CAV1 WT
CAV1 WT
PACSIN2
E
CAV1 WT
PACSIN2
CAV1 WT
PACSIN2
F
Myc : PACSIN2
Pearson's Correlation
***
***
ns
CAV1 WT myc + - - +
CAV1 WT mCherry - - + +
P132L myc - + + -
G
CAV1 mCherry : PACSIN2
Pearson's Correlation
ns
ns
ns
CAV1 WT myc - + -
CAV1 WT mCherry + + +
P132L myc - - +
H
Myc : CAV1-mCherry
Pearson's Correlation
*
CAV1 WT myc + -
CAV1 WT mCherry + +
P132L myc - +

**Figure S5. When co-expressed with WT CAV1, P132L partially co-localizes with endogenous PACSIN2.** CAV1 $^{-/-}$ MEFs were transiently transfected with **(A)** CAV1-mCherry, **(B)** P132L, **(C)** CAV1-mCherry plus P132L, **(D)** CAV1-mCherry plus CAV1, or **(E)** CAV1 for 24h, fixed, and immunostained prior to imaging. Representative AiryScan images are shown. Bar, 5 µm. **(F-H)** Pearson's correlation analysis was carried out using ImageJ (n = 90 ROIs from 3 independent experiments). A one-way ANOVA with Tukey's test (≥3 groups) or an unpaired Student's t test (2 groups) was used to calculate P-values. n.s., not significant.

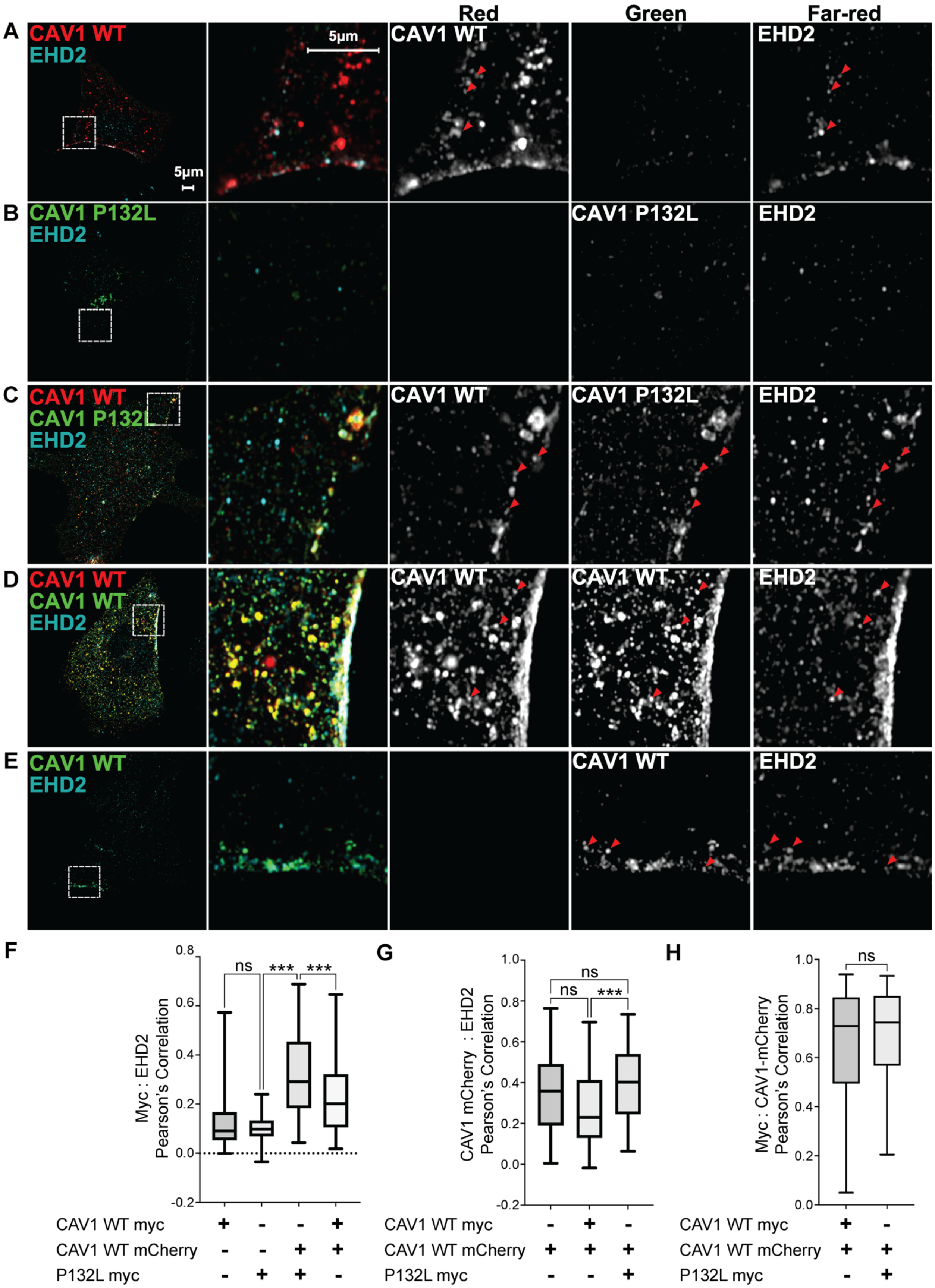

Red
Green
Far-red
A
CAV1 WT
EHD2
5µm
5µm
CAV1 WT
EHD2
B
CAV1 P132L
EHD2
CAV1 P132L
EHD2
C
CAV1 WT
CAV1 P132L
EHD2
CAV1 WT
CAV1 P132L
EHD2
D
CAV1 WT
CAV1 WT
EHD2
CAV1 WT
CAV1 WT
EHD2
E
CAV1 WT
EHD2
CAV1 WT
EHD2
F
Myc : EHD2
Pearson's Correlation
0.8
0.6
0.4
0.2
0.0
-0.2
ns
***
***
CAV1 WT myc + - - +
CAV1 WT mCherry - - + +
P132L myc - + + -
G
CAV1 mCherry : EHD2
Pearson's Correlation
1.0
0.8
0.6
0.4
0.2
0.0
-0.2
ns
ns
***
CAV1 WT myc - + -
CAV1 WT mCherry + + +
P132L myc - - +
H
Myc : CAV1-mCherry
Pearson's Correlation
1.0
0.8
0.6
0.4
0.2
0.0
ns
CAV1 WT myc + -
CAV1 WT mCherry + +
P132L myc - +

**Figure S6. P132L partially colocalizes with endogenous EHD2 when co-expressed with WT CAV1.** CAV1 $^{-/-}$ MEFs were transiently transfected with **(A)** CAV1-mCherry, **(B)** P132L, **(C)** CAV1-mCherry plus P132L, **(D)** CAV1-mCherry plus CAV1, or **(E)** CAV1 for 24h, fixed, and immunostained prior to imaging. Representative AiryScan images are shown. Bar, 5 μm. **(F-H)** Pearson's correlation analysis was carried out using ImageJ (n = 90 ROIs from 3 independent experiments). A one-way ANOVA with Tukey's test (≥3 groups) or an unpaired Student's t test (2 groups) was used to calculate P-values. n.s., not significant.
